# A Novel Drug Candidate that Selectively Targets the Critical Androgen Receptor-ELK1 Growth Axis in Advanced and Drug-Resistant Prostate Cancer

**DOI:** 10.64898/2026.05.26.727474

**Authors:** Claire Soave, Lisa Polin, Charles Ducker, Vy Ong, Seongho Kim, Luke Pardy, Jing Li, Xun Bao, Yanfang Huang, Peter E. Shaw, Rahul Khupse, Manohar Ratnam

## Abstract

Androgen receptor (AR)-dependent prostate cancer (PCa) cells require co-activation of ELK1 by AR to activate a critical set of cell cycle and mitosis genes, regardless of hormone - sensitivity. A small molecule antagonist (KCI807) that inhibits AR-dependent growth by selectively binding to AR and blocking its association with ELK1 is limited as a drug by auto-induced metabolism. Using structure-activity data, consistent with modeling a physically mapped KCI807 binding pocket, we developed a new class of compounds with a different core structure comprising 5-Hydroxy-2-(3-hydroxyphenyl)-1-methylquinolin-4(1H)-one (KCI830), with variable N- substituents. The compound with a N-2,2,2-trifluoroethyl substitution (KCI838) was the fastest acting and most potent inhibitor of AR-dependent cell growth and colony formation in PCa model cells, including exclusively AR splice variant-dependent and other enzalutamide-resistant cells, without affecting growth of AR-negative cell lines. Critical tests were conducted to establish that KCI838 recapitulates the previously elucidated mode of action of KCI807. KCI838 selectively inhibited ELK1-dependent vs. androgen response element (ARE)-driven promoter and gene activation by AR. KCI838 blocked AR binding to ELK1 *in situ* tested by BRET assay. Increasing the total cellular AR by ∼2-fold using ectopic AR expression caused the predicted change in drug dose-response profile for growth, implicating AR as the exclusive target for the activity of KCI838. KCI838’s molecular scaffold conferred reduced enzyme induction in primary human hepatocytes and weakened interactions with human UGT1A1 and CYP1A2. In mice bearing an aggressive, enzalutamide-resistant patient-derived PCa tumor xenograft characteristically overexpressing prostatic acid phosphatase, daily bolus injections of a soluble 3’phosphate monoester prodrug of KCI838 (KCI838PME) progressively inhibited tumor growth, concomitant with tumor accumulation of active hydrophobic drug, without significant toxicity. Additionally, ALZET osmotic pumps were used to establish proof-of-concept for reversible *in vivo* anti-tumor activity of KCI838PME administered in a low dose, controlled release mode. The results warrant investigation of KCI838PME in a controlled-release formulation, to treat PCa that is resistant to current AR-targeted therapies while obviating the need for testosterone suppression.

## Introduction

Prostate cancer (PCa), whether early stage or advanced, is typically dependent on AR for growth (1, 2). Advanced PCa is commonly treated by androgen deprivation therapy (ADT), via chemical castration (3, 4) as well as treatment with high affinity androgen antagonists (enzalutamide, apalutamide and darolutamide) (5) or an inhibitor of intratumor testosterone synthesis (e.g., abiraterone) (6). Unfortunately, the patients frequently develop castration-resistant PCa (CRPC) which could also become resistant to androgen antagonists and abiraterone. The resistant tumors nevertheless continue to depend on AR, whose growth signaling role may be restored, largely through amplification of AR (7) or expression of its splice variants (AR-Vs), which lack the ligand binding domain (8–10). Splice variants of AR are expressed in ∼90 percent of metastatic tumors and may work either in concert with or independent of full-length AR (11–16). Resistance mechanisms also include hormone-independent crosstalk between AR and certain signaling pathways, alterations in the AR co-regulator complement or mutation of AR (7). ADT also affects systemic non-growth-related functions of AR, resulting in adverse side effects that are both acute (fatigue, hot flashes), long-term (hyperlipidemia, insulin resistance, cardiovascular disease, anemia, osteoporosis, sexual dysfunction and cognitive defects) and include loss of the feeling of well-being (3, 17, 18).

As summarized in this section, we have previously preclinically validated a new strategic approach that addresses the dual limitation of ADT by disrupting a functional arm of AR that is (i) preserved as a crucial mechanism for supporting growth in CRPC, including enzalutamide resistant cancer, and (ii) only necessary for tumor growth stimulation by AR but not for other gene regulatory functions of AR that are unrelated to growth. This new drug target is a complex of AR with the ETS family transcription factor ELK1.

ELK1 is a downstream effector of the MAPK signaling pathway and belongs to the ternary complex factor (TCF) sub-family of the ETS family of transcription factors. ELK1 characteristically binds to purine-rich GGA core sequences (19) and is in a repressive or passive association with many cell proliferation genes. Phosphorylation by ERK transiently hyper-stimulates ELK1 to activate its target genes including association of ELK1 with serum response factor (SRF) for activation of immediate early genes (19–21). Chromatin sites of AR binding are highly enriched for ELK1 binding DNA *cis*-elements (22). Transcriptome analysis of PCa cells, following individual and combined knockdown of AR and ELK1, showed that ELK1 is fully or partially required for a substantial proportion (∼ 27 percent) of all gene activation by androgen in PCa cells and that this gene subset is primarily enriched for growth functions (23). ELK1 was not required for regulation of non-growth-related genes by AR (23, 24). Consistent with this finding, depletion of ELK1 in a variety of hormone-dependent or -independent AR-positive PCa cell lines completely inhibited growth, colony formation and tumor formation ( 23); in contrast ELK1 depletion did not affect AR-negative PCa cell lines or other cell types that did not express AR (23).

ELK1 interacts with AR in situ by anchoring AR to chromatin sites of ELK1 binding (23, 24). This enables constitutive activation of target (growth) genes by AR. This coactivator role of AR is not associated with ELK1 phosphorylation or MAPK signaling and does not require the transactivation function of ELK1 (23, 24). AR binds to ELK1 (Kd = 2 × 10^-8^ M) by utilizing the two ERK docking sites in ELK1, largely through the amino-terminal A/B domain of AR which lacks the ligand binding site (24). AR binding to ELK1 was essential for growth, as demonstrated by the dominant-negative effect of an AR docking site mutant of ELK1 on growth of PCa cells that are insensitive to MEK inhibition (24). Within AR, the sites of interaction with ELK1 have been mapped to two peptide segments within amino-terminal domain (NTD) spanning residues 358-457 and 514-557 that were both required for PCa cell growth (25). The major splice variants of AR also supported ELK1-dependent gene activation (23, 24). ELK1 is also a strong and independent prognosticator of PCa recurrence (26). Therefore, disruption of the ELK1-AR complex using a suitable AR-binding molecule should selectively suppress growth of PCa/CRPC tumors, including tumors dependent on AR splice variants, without significant effects on other functions of testosterone/AR and without affecting the normal functions of ELK1. Proof of this concept was provided through the discovery of a small molecule inhibitor of the binding of AR to ELK1, KCI807 (27).

The discovery and detailed mechanism of action of KCI807 in PCa cells has been reported previously and defined by a narrow set of structural and conformational requirements (27, 28). KCI807 binds to AR with a Kd of 7 × 10^-8^ M, i.e., with about a third of the binding affinity of ELK1 for AR (27). Binding of KCI807 to AR blocks the association of ELK1 in a dose-dependent manner, demonstrable both in cell-free systems and *in situ* (27). KCI807 selectively prevents recruitment of AR to chromatin sites by ELK1 without interfering with the binding of AR to canonical androgen response elements (AREs) in chromatin (27). KCI807 also inhibits ELK1-dependent promoter activation by the major AR splice variant AR-V7 and mutant AR that are resistant to enzalutamide (27). Although KCI807 inhibits constitutive co-activation of ELK1 by AR, it does not interfere with the transient activation of ELK1 by ERK (27). Based on transcriptome analysis in PCa cells, the target gene set of KCI807 was principally associated with functional roles in cell cycle progression and mitosis within the AR signaling axis, and virtually exclusively associated with synergistic activation by ELK1 and AR (27). The cell growth inhibitory effect of KCI807 was selective for PCa cells that were dependent on AR and/or AR-V7 or enzalutamide-resistant mutant AR with a better growth inhibitory profile than enzalutamide in well-established prostate cancer cell line models (27). Evidence supports a mechanism of action of KCI807 in which the molecule binds to a putative binding pocket within the DNA binding domain (DBD) of AR outside the DNA contact surface of the DBD, inducing a conformational change that occludes ELK1 binding at adjacent sites within the NTD (28).

KCI807 inhibited AR-positive CRPC tumor growth *in vivo,* including aggressively growing, enzalutamide-resistant and AR-V7 positive tumors and also aggressively growing enzalutamide-resistant patient-derived (PDX) CRPC tumors (27). However, the compound dose required for its antitumor effect was relatively high (250 mg/kg administered i.p. on alternate days), and tumor growth eventually resumed, coincident with a striking decline in plasma levels of the compound (27). The dramatic decline in its plasma levels after administering the compound for a sustained period was clearly due to its self-induced metabolism in the rodent model. The major metabolic products of KCI807 resulted from oxidation at the C5’ position of a 3’-hydroxyphenyl ring, as well as conjugation of its 3’OH group as a glucuronide, glycosylate and sulfate, (27).

To overcome the above limitations of KCI807, we designed a new class of small molecules (the KCI830 series) as potential drug candidates to treat PCa resistant to castration, enzalutamide and abiraterone. This class of molecules was rationally designed using a large body of structure-activity studies (SAR) combined with modeling of the KCI807 binding within a physically mapped putative binding pocket within AR (SAR data and related modeling studies will be published elsewhere). Here we report on the biological characteristics of KCI838, which showed the most promise among them, with greater potency and metabolic viability than KCI807. The study design includes critical tests to establish that KCI838 recapitulates the biological activity and target selectivity of KCI807 but with characteristics more suitable for clinical development.

## Materials and Methods

### Reagents

Rabbit monoclonal antibody to AR (ab133273, 1:500) was bought from Abcam. Antibody to GAPDH (sc-47724, 1:3000) was bought from Santa Cruz Biotechnology. Rabbit monoclonal antibody to ELK1 (ab32106, 1:1000) was purchased from Abcam. Mouse monoclonal antibody to β-actin (ab6276, 1:5000) was bought from Abcam. Testosterone was from Sigma-Aldrich. Dihydrotestosterone was from Sigma-Aldrich. R1881 was a kind gift received from Stephan Patrick, Wayne State University (Detroit, MI). Enzalutamide was from Selleck Chemicals. Chemicals and reagents needed for synthesis of the KCI830 series compounds were all purchased from Sigma-Aldrich. Lipofectamine ™ 2000 was from Thermo Scientific. KCI807 was from INDOFINE Chemical Company. Plasmids pLVX-AR-V7, pLVX control, RLuc8.6-AR and Turbo635FP-AR kindly shared by Dr.Yan Dong, Tulane University (New Orleans, LA). Turbo-ELK1 was generated by replacing AR coding sequences in Turbo635FP-AR with coding sequences for ELK1.

### Synthesis of N-substituted 5,3’-dihydroxyphenylquinolone compounds

Compound design and synthetic procedures comprise intellectual property of Wayne State University, covered by patent #US12,458,636 B2. Scale up syntheses and purification were outsourced to a commercial custom synthesis company. The new method of chemical synthesis was carried out per the schematic showed in **Supplemental Figure 1** using the procedures described below.

#### Synthesis Step 1

1mmol of flavone is suspended in 25 mL of trimethyl orthoformate or triethyl orthoformate. Then 1 mL of concentrated sulfuric acid (98%) is added slowly at room temperature, with evolution of gas. The mixture is stirred for 12 hours at room temperature in a closed system attached to a bubbler. Then, 20 mL of ethyl acetate is added to the mixture at room temperature. The precipitated solids were filtered off, washed with 5 mL ethyl acetate and dried in air for 2 hours. The crude filtered solid, yellow sulfate salt was used in the next step without purification.

#### Synthesis Step 2

The intermediate from step 1 (0.5 mmol) is weighed in a flask and suspended in 2 mL of neat amine when the color changes to orange. The mixture is stirred overnight. The progress of the reaction is monitored by thin layer chromatography (Dichloromethane: Methanol 10:1). The solvent is evaporated and the crude solid is purified by precipitation using 12 mL acetone: hexane (5:1 ratio) or by column chromatography. The precipitated product is filtered off dried and spectral analysis is carried out. NMR (Nuclear Magnetic Resonance, ^1^H, ^13^C and ^19^F), mass spectrometry and CHN elemental analysis are used to confirm the structures of the purified products. The amine serves as both reactant and solvent and the amine compound used is based on the R1 group desired on nitrogen. For example, 2,2,2-trifluoroethylamine was used when the R1 group was trifluoroethyl. When the R1 group does not contain fluorine, the crude products are recrystallized from hot anhydrous ethanol. For fluorinated derivatives, alumina is used for initial small scale purification using hexane: isopropanol (4:1 ratio).

### Synthesis and purification of disodium salt of KCI838 3’-phosphate monoester (KCI838PME)

Synthesis was carried out per the schematic showed in **Supplemental Figure 2** using the procedures described below:

For synthesis of **3**, 6.5mmol of KCI838 was suspended in 30 mL of TEA and 30 mL of CCl4 at 0 degree C, followed by 7.2 mmol of triethyl phosphite added dropwise. The mixture was warmed to room temperature overnight. After workup, compound **3** was purified by silica chromatography to give compound **3** as an oil. For synthesis of **4**, 4.5 mmol of compound **3** was dissolved in 50 mL of DCM, to which 42 mmol of TMSBr was added. The mixture was stirred overnight at room temperature and then concentrated to remove excessive TMSBr in vacuo. The residue was coevaporated with toluene for 3 times. The residue was triturated with water and filtered to give **4** as an orange solid. The solid was further washed with cold MeCN and ether, and dried. For synthesis of **5**, 3.8 mmol of compound **2** was dissolved in MeOH and 7.7 mmol of NaOMe was added at room temperature. The mixture was stirred for 15min. MeOH was removed *in vacuo* and the residue was purified by HPLC.

The preparative HPLC purification of KCI838PME (compound **5, Supplemental Figure 2**) was performed on Waters 2545 preparative HPLC. Vydac prep C18 HPLC column (50*250mm 10-Micron) was used for preparation. Gradient: pH 8-9 NaOH in water as mobile phase A and acetonitrile as mobile phase B were used. The flow rate is 75 mL/min. The gradient starts as 5% B and holds for 3 minutes, increases from 5% B to 95 % B in 18 minutes, and hold for 6 minutes, then go back to 5% B in 3 minutes. Pure fractions were collected, evaporated and dried to afford the pure compound as red solid. 613 mg of crude product in 15.5 mL of water was purified to afford about 220 mg of the desired compound.

### Cell lines

22Rv1 cells, LNCaP cells, VCaP cells, DU145 cells, A549 cells, MDA-231 cells and HeLa cells were purchased from ATCC and frozen after less than 5 passages. The cells were therefore not further authenticated or tested for mycoplasma. The CRISPR-generated CWR22Rv1-AR-EK cells (16) which lack full-length AR and are dependent on endogenous AR splice variants (AR-Vs) were purchased from Cancer Research Technology Limited (London, UK). All cells underwent <10 passages during the studies. The growth media used were as follows. For 22Rv1 cells, RPMI 1640 (phenol red-free), 10% FBS, 1% penicillin/streptomycin, 1% L-glutamine. For VCaP cells, DMEM high glucose (phenol red-free), 10% FBS, 1% penicillin/streptomycin, 1% L-glutamine, 1 nM dihydrotestosterone (DHT). For LNCaP cells, RPMI 1640 (phenol red-free), 10% FBS, 1% penicillin/streptomycin, 1% L-glutamine, 1% sodium pyruvate, 1% HEPES, 0.1 nM R1881. For DU145 cells, DMEM high glucose (phenol red-free), 10% FBS, 1% penicillin/streptomycin, 1% L-glutamine. For MDA-MB-231 cells, DMEM high glucose (phenol red-free), 10% FBS, 1% penicillin/streptomycin, 1% L-glutamine. For A549 cells, RPMI 1640 (phenol red-free), 10% FBS, 1% penicillin/streptomycin, 1% L-glutamine. For HeLa cells, DMEM high glucose (phenol red-free), 10% FBS, 1% penicillin/streptomycin, 1% L-glutamine.

### Recombinant HeLa Cells

The development of the two types of recombinant HeLa cells used in this study have been previously described (27). One of the recombinants has a stably integrated minimal promoter-luciferase reporter containing five upstream Gal4 elements (Gal4-TATA-Luc) and also constitutively expresses a Gal4-ELK1fusion protein in which the Gal4 DNA binding domain is substituted for the ETS DNA binding domain of ELK1. These cells further stably express recombinant full-length human AR. The second type of recombinant HeLa cells have a stably integrated minimal promoter-luciferase reporter and an upstream androgen response element (ARE) sequence. These cells also stably express recombinant full-length human AR.

### Cell Viability Assay

Cells from monolayer cultures were trypsinized and seeded in 96-well plates coated with poly(d-lysine) at a density of 4000 cells per well (for 22Rv1 cells and LNCaP cells) or 5000 cells per well (for VCaP cells). The media used were the same as the growth media described above for each cell line. The plates were incubated at 37°C in 5% CO2. Twenty-four hours later, the cells were treated with the indicated compound or dimethylsulfoxide (vehicle). The culture medium was replaced with fresh media containing vehicle or the appropriate concentration of compound at the end of 48h during the time course of the assay. Cell viability was determined on Day 0 and at the various time points using the standard 3-(4,5-dimethylthiazol-2-yl)-2,5-diphenyltetrazolium bromide (MTT) assay. The assays ware conducted in replicates of six wells per condition and values plotted as percent of the Day 0 values.

### Colony formation assay

22Rv1 cells were plated in triplicate at a density of 2000 – 6000 cells/well in poly-D lysine-coated 6-well plates in 48-hour conditioned medium (phenol-red free RPMI1640 10% FBS, 1% penicillin/streptomycin, 1% L-glutamine) obtained from monolayer cultures of the same cells, as previously described. 48h after plating, the cells were treated with various concentrations of the test compounds in the same media. In certain experiments, 1nM dihydrotestosterone (DHT) was included in the growth media. The test compounds and media were replenished every 48 hours until colonies developed in the control untreated wells (in 7 - 10 days). The colonies were then stained with crystal violet and counted using the GelCount colony counter (Oxford Optronix). The IC50 and IC75 values for inhibition of colony formation were calculated using GraphPad Prism 5 software.

### RNA Isolation, Reverse Transcription, and Quantitative Real Time PCR

Total RNA from cells was isolated using the PureLink RNA Mini Kit (Invitrogen, ThermoFisher Scientific) according to the manufacturer’s protocol. Reverse transcription was performed using 30 ng of total RNA and the High Capacity cDNA Reverse Transcription kit (Applied Biosystems, ThermoFisher Scientific) according to the vendor’s protocol. cDNA was measured by quantitative real time PCR using the StepOnePlus Real-Time PCR System (Applied Biosystems, Invitrogen) and TaqMan Fast Universal PCR Master Mix (Applied Biosystems, ThermoFisher Scientific). All primers and TaqMan probes were purchased from the Applied Biosystems inventory (ThermoFisher Scientific). All samples were measured in triplicate and normalized to the values for *GAPDH*.

### Western blots

Cells were lysed for western blot sample preparation using radioimmune precipitation assay buffer (150 mM NaCl, 1% Nonidet P-40, 0.5% sodium deoxycholate, 0.1% SDS, 50 mM Tris of pH 8.0) which contained a protease inhibitor mixture and EDTA (78410, Thermo Fisher Scientific), then incubated on ice for one hour. Protein concentration was measured using a Bradford assay (500006, Bio-Rad). Lysed samples were heated for 5 min at 95°C, and then proteins (2-5 µg) in the samples were separated by electrophoresis on 8% SDS-polyacrylamide gels and electrophoretically transferred to PVDF membranes (IPVH00010, Millipore). The membranes were probed with primary antibody and secondary horseradish peroxidase-conjugated antibody. Proteins were visualized using the WesternBright ECL Spray (K-12049-D50, Advansta) and a ChemiDoc Touch Imaging System (Bio-Rad).

### Hepatic enzyme induction

Primary human hepatocytes used were the “Cryo Human Hepatocytes” from Gibco. 24 hours prior to treatment, cells were plated at 350,000 cells/well (700,000 cells/mL) in the following media: Williams E medium (WEM, A1217601), 2.5 mL FBS, 5 µL 10 mM dexamethasone (from Gibco Thawing/Plating Supplement Pack, CM3000), 1.8 mL thawing/plating cocktail (from Gibco Thawing/Plating Supplement Pack, CM3000). 5 hours after plating, media was replaced with the following media: Williams E medium (WEM, A1217601), 0.5 µL 10 mM dexamethasone (from Gibco Cell Maintenance Supplement Pack, CM4000), 2 mL cell maintenance cocktail (from Gibco Cell Maintenance Supplement Pack, CM4000). Treatment media was same as cell maintenance media listed above. Cells were treated with the control inducers (b-naphthoflavone, phenobarbital, dexamethasone and rifampin), KCI807 (0.5uM, 5uM and 50uM) or KCI838 (0.5uM, 5uM and 50uM), incubated at 37 degrees in 5% CO2 for 72h and then harvested for RNA extraction. The mRNAs for the following hepatic enzymes were quantified as described above. Monooxygenases: CYP1A2, CYP2A6, CYP2B6, CYP2C9, CYP2C19, CYP3A4; Sulfotransferases: SULT1A1, SULT2A1; UDP-glucuronyltransferases: UGT1A1, UGT1A4, UGT1A6, UGT1A9, UGT2B4; UDP-glycosyltransferase: UGT2B7.

### CYP1A2 inhibition assay

This assay was conducted by the specialized service company, Cyprotex US LLC (Watertown, MA). A stock solution of KCI838 was prepared in DMSO at 50 mM and stored at −20 °C. Serial dilutions of the stock solution were prepared in acetonitrile:DMSO (9:1) for CYP inhibition testing. The final DMSO content in the reaction mixture was equal in all solutions used within an assay and was ≤ 0.2 %. KCI838 was incubated at seven increasing concentrations with pooled human liver microsomes (Bioreclamation-IVT, Baltimore MD) in the presence of 2 mM NADPH in 100 mM potassium phosphate (pH 7.4) containing 5 mM magnesium chloride and the probe substrate, tacrine (5uM), in a 200 μL assay (final volume). The selective CYP1A2 inhibitor α-naphthoflavone was screened alongside KCI838 as a positive control. After incubation for 10 min at 37 °C (**Table 1**), the reactions were terminated by addition of methanol containing internal standard for analytical quantification. The quenched samples were incubated at 4° C for 10 min and centrifuged at 4 °C for 10 min. The supernatant was removed and the probe substrate metabolite analyzed by LC-MS/MS. A decrease in the formation of the metabolite compared to vehicle control was used to calculate an IC50 value (the test concentration that produces 50% inhibition).

**Table 1:** Mouse weights during daily bolus injections of KCI838PME.

|  | Day<br>1 | Day<br>2 | Day<br>3 | Day<br>4 | Day<br>5 | Day<br>6 | Day<br>7 | Day<br>8 |
| --- | --- | --- | --- | --- | --- | --- | --- | --- |
| Control<br>Group (g) | 20.1 | 20.3 | 20.6 | 21 | 21 | 21 | 21.4 | 21.5 |
| Treatment<br>Group (g) | 20.4 | 20 | 20 | 20 | 20 | 20 | 20 | 20.3 |

### UGT1A1 inhibition assay

This assay was conducted by the specialized servive company, Cyprotex US LLC (Watertown, MA). KCI838 was incubated at seven increasing concentrations with human UGT1A1-expressed SupersomesTM (0.25 mg/mL), alamethicin (25 μg/mL) and UDPGA (5 mM) in the presence of the probe substrate estradiol (10 μM) for 30 min at 37 °C. The UGT1A1 inhibitor, atazanavir, was screened alongside the test compounds as a positive control. The reactions were terminated by quenching with one volume of methanol containing an analytical internal standard. The samples will be centrifuged at 5000 rpm for 10 min at 4 °C. The metabolites were monitored by LC-MS/MS and a decrease in the formation of the metabolite compared to the vehicle control was used to calculate an IC50 value (test compound concentration which produces 50 % inhibition).

### Promoter inhibition assays

To measure ELK1-dependent promoter activation by AR, we used recombinant HeLa cells, which have a stably integrated minimal promoter -luciferase reporter containing five upstream Gal4 elements (Gal4-TATA-Luc) and constitutively express a GAL4-ELK1 fusion protein (in which the Gal4 DNA binding domain is substituted for the ETS DNA binding domain of ELK1), as well as full-length AR. To measure androgen response element (ARE)-driven promoter activation by AR, we used recombinant HeLa cells harboring a minimal promoter-luciferase reporter with an upstream ARE sequence (ARE-TATA-Luc) and also stably expressing full-length AR. The two types of recombinant cells were grown in DMEM supplemented with 5% FBS or 10% FBS, respectively, and 100 units/mL penicillin, 100 µg/mL streptomycin, 2mM L-glutamine mixture (Invitrogen). The selection antibiotics used in maintenance cultures of the two recombinant cell types were 100 µg/mL Hygromycin (Invitrogen) (to maintain GAL4-ELK1), 100 µg/mL Geneticin (Invitrogen) (to maintain Gal4-TATA-Luc) and 2 µg/mL Puromycin (Sigma-Aldrich) (to maintain AR) and 400 ug/mL Geneticin (Invitrogen) (to maintain ARE-TATA-Luc). The cells were hormone-depleted 24 hours prior to screening in media containing heat-inactivated, charcoal-stripped serum. Cells were then plated in 96-well white flat bottom plates (20,000 cells/well) (Corning product #3917) and incubated for approximately 18 hours prior to treatment. The following day, cells were treated with the appropriate doses of test compounds along with testosterone or dihydrotestosterone (DHT) (final concentration, 1 nM), using 6 replicate wells per dose. Vehicle controls (DMSO, 0.1% v/v) were included. The plates were incubated for 6 hours at 37°C in 5% CO2. The media was then aspirated and 20 µL of Promega Glo Lysis Buffer (Promega #E2661) was added to each well, and then the plates were incubated at room temperature for 10-15 minutes on an orbital shaker. Luciferase activity was measured for each well using firefly luciferase substrate from the Luciferase Assay System (Promega) in a luminometer (Centro XS3 LB 960, Berthold, Wildbad, Germany).

### AR target gene expression assays

22Rv1 cells were grown in RPMI supplemented with 10% FBS and 100 units/mL penicillin, 100 µg/mL streptomycin, 2mM L-glutamine mixture (Invitrogen). The day before treatment, cells were plated in poly-D lysine coated plates (100,000 cells/well) (Corning #354461) in phenol red-free growth media. The following day, cells were treated with 10 µM of each compound or DMSO vehicle (0.1% v/v). Cells were incubated for 72 hours at 37°C in 5% CO2, with treated media replenished at 48 hours. At 72 hours, total RNA from cells was isolated using the PureLink RNA Mini Kit (Ambion, Invitrogen) according to the manufacturer’s protocol. Reverse transcription was performed using 250 ng of total RNA and the High Capacity cDNA Reverse Transcription Kit (Applied Biosystems, Invitrogen) according to the manufacturer’s protocol. cDNA was measured by quantitative real time PCR using the StepOnePlus Real-Time PCR System (Applied Biosystems, Invitrogen) and TaqMan Fast Universal PCR Master Mix (Applied Biosystems, Invitrogen). All primers and TaqMan probes were purchased from the Applied Biosystems inventory (Invitrogen). Samples were measured in triplicate and normalized to the values for GAPDH.

### BRET assay for inhibition of in situ binding of AR to ELK1 by KCI838

HEK293T cells were seeded in 6-well dishes at 2×10^5^ cells/ml, and 24 hours later were transfected with Rluc-AR alone, and alongside Turbo-ELK at a 4:1 ratio of ELK1:AR using Mirus LT1 reagent (The 4:1 ratio was determined to be the optimal ratio for the inhibition assay after testing different ratios of Turbo - ELK: Rluc-AR). After 24 hours, cells were treated with DMSO (vehicle control) or KCI838 (0.4uM and 10uM) and left for 24 hours. They were then trypsinized and seeded in 96 well white-bottomed plates at 100,000 per well (in quadruplicate) in media lacking phenol red (with DMSO or drug as required), left for 2 hours to settle, and then treated with 25uM coelenterazine and luminescence read at 535 and 635nm on a Clariostar (BMG Labtech) dual plate reader. BRET readings were calculated as 635/535 ratio minus 635/535 ratio for Rluc-AR only transfected controls. Cell lysates were probed by western blot for AR or ELK1, probing for β-actin as the loading control. Cell viability in the presence of KCI838 was ensured by treating HEK293T cells (3×10^5^ cells/ml seeded in 6-well dishes 24 hours pre-treatment) for an extended period (3 days) with the compound or vehicle control and measuring viability using trypan blue staining.

### Generation of recombinant 22Rv1 cells overexpressing AR

AR expressing lentiviral expression plasmid pCDH-CMV-MCS-EF1-puro (System BioSciences) or the plasmid vector alone (control), was packaged in 293FT cells. Virus-containing media was harvested at 48 and 72 hours after packaging and stored at −80°C. 48 hours before infection, 22Rv1 cells were plated at 30-50% confluence in 10 cm dishes in phenol-red free medium supplemented with 10% heat-inactivated FBS and 2 mM L-glutamine. On the day of infection, cells were infected with the lentivirus with polybrene (8 μg/mL) for a duration of 5 hours. The virus was then replaced with fresh medium containing 10% FBS and 100 units/mL penicillin, 100 µg/mL streptomycin, and 2mM L-glutamine mixture. Cells were placed under puromycin selection for at least 4 passages, at a previously optimized concentration of 0.4 ug/mL puromycin to select for the pools of transduced cells.

### Immunohistochemistry

Immunohistochemistry staining of paraffin embedded sections human prostate tumor tissue (PDX) samples was conducted by iHisto Inc. Slides were loaded into a LeicaBond RX with covertiles and stained as follows: Slides were baked and deparaffinized (Bond Dewax Solution). Antigen retrieval was performed. The slides were brought to 95 degrees Celsius for 20 minutes while incubating in Bond Epitope Retrieval 2 (pH 9). Slides were washed 3 times for 2 minutes each (Bond Wash Solution). Slides were blocked with 2.5% normal horse serum for 20 minutes at room temperature. Slides were rinsed. Primary antibody diluted in 2.5% horse serum Alk Phsp Nonspecific ab 249-MSM2-P0 1:100, and PSAP 55-MSM3-P0 1:100 was applied to the sections and allowed to incubate for 30 minutes at room temperature. Slides were washed 3 times for 2 minutes each (Bond Wash Solution). Secondary antibody HRP horse anti-mouse was applied to the sections and allowed to incubate for 8 minutes at room temperature. Slides were washed 3 times for 2 minutes each (Bond Wash Solution). DAB was applied to the sections and allowed to stain for 5 min for Alk Phsp Nonspecific ab 249-MSM2-P0 and 3 min for PSAP 55-MSM3-P0. Slides were washed 3 times for 5 minutes each (Bond Wash Solution). Sections were stained with hematoxylin for 35 seconds at room temperature. Slides were rinsed with deionized water twice for 2 minutes each. Once automated staining in the Leica Bond RX was completed, slides were removed to dehydration system 100% ethanol 3x, for 5 minutes each time and 5 minutes on xylene then coverslipped.

### In vivo studies

The patient-derived prostate tumor xenograft model (PDX-PR011) has been previously described (27). This tumor was derived from a bone biopsy from a patient with CRPC at Karmanos Cancer Institute. The tumor was implanted SC into a SCID male mouse, where it metastasized to the lungs. The metastatic lungs were harvested and implanted into a fresh mouse, where it formed a SC tumor. The tumor xenograft retained AR expression (27).

For the drug efficacy studies using bolus injections of the drug, the tumor was implanted sc, bilaterally in groups of 7 mice. The treated group was injected daily on Days 1-8 with KCI838PME. The bolus injections were administered iv at a drug dose of 4mg per mouse per day in water. Treatment with KCI838PME was initiated within 1 day of tumor implantation because of the relatively highly aggressive growth rate of the tumors (8 days to reach tumor growth to about 1g in the placebo control group). Tumor measurements were conducted using a caliper and tumor masses (in mg) estimated by the formula, mg = (a × b2)/2, where “a” and “b” are tumor length and width in mm, respectively. The experiment was terminated when cumulative tumor burdens reached 5% to 10% of body weight (1–2 g) in the control group. Six hours after the 8^th^ bolus injection, the mice were sacrificed after drawing blood samples for preparation of plasma and measurement of plasma levels of KCI838PME and KCI838. At the same time, the residual tumors were harvested to measure tumor levels of KCI838PME and KCI838. In a separate experiment, 9 non-tumor bearing mice were injected 180 mg/kg of KCI838PME iv in water and blood samples drawn from groups of 3 mice at 0.5h, 2h and 6h for preparation of plasma and measurement of plasma levels of KCI838PME and KCI838.

For studies using controlled release of KCI838PME, on Day 0, the tumor was implanted sc, bilaterally in groups of 7 mice. On Day 1, the mice were implanted with two ALZET pumps per mouse (1-week release, 200 ul capacity) containing either water (control group) or a 20mg/ml solution of KCI838PME in water (total, 8mg KCI838PME per mouse in the treatment group). The ALZET pumps were primed overnight before implantation, following the manufacturer’s directions. The drug was exhausted on Day 8 but all mice were kept until Day 14. Tumor volumes were measured as described above, on Days 7, 10, 12 and 14. In a separate experiment, groups of 3 non-tumor bearing mice were implanted primed ALZET pumps (2 pumps per mouse, containing a total of 8 mg KCI838PME in water, per mouse). Two days and 7 days after implantation of the pumps, plasma was obtained from the mice to measure levels of KCI838PME and KCI838.

Mouse weight and behavior were monitored daily in all of the above studies.

### Assay of KCI838 and KCI838PME in plasma and tumor tissue samples

All LC-MS/MS analyses were carried out on a Waters Xevo TQ-XS LC-MS/MS mass-spectrometer coupled with a Waters Acquity H-class UPLC system (Milford, MA). The instrument operation and data acquisition were under control by MassLynx® 4.2 software, and data processing and quantification were carried out with TargetLynx® XS software.

Chromatographic separation was optimized on a Waters Xbridge C18 column (3.5 µm, 2.1 x 50 mm) with a gradient elution consisting of mobile phase A (0.1% formic acid in water) and mobile phase B (0.1% formic acid in methanol) at a flow rate of 0.3 mL/min. The elution gradient program was as follows [shown as the time (min), (% mobile phase B)]: 0, 10%; 0.5, 10%; 1.5, 90%; 3.0, 90%; 3.5, 100%; 4.5, 100%; 4.7, 10%; 7, 10%. The retention times of KCI838 and KCI838 PME are 2.8 min and 2.6 min, respectively. The column was maintained at 35 °C. A mixture of methanol and water (75/25, v/v) was used as needle wash to minimize the carryover. The Waters Xevo TQ-XS mass spectrometer was operated in electrospray positive ionization using multiple reaction monitoring mode (MRM). The capillary voltage was set at 3.90 kV, and the desolvation temperature 250 °C. Gas flow parameters were optimized to 600 L/Hr for desolvation, 150 L/Hr for cone, and 7.0 Bar for nebulizer, respectively. The dwell time was set at 100 ms. Cone voltage and collision energy were optimized at 66 V and 42 eV for KCI838, and 2 V and 46 eV for KCI838 PME. The most sensitive MS transition of m/z 336.12 → 210.18 and 416.16 → 307.19 were selected for monitoring KCI838 and KCI838 PME, respectively.

KCI838 and KCI838 PME stock solutions were prepared in methanol at a final concentration of 5 mM and stored in brown glass vials at – 80 °C. Their corresponding working solutions were prepared separately and freshly by serial dilutions of the stock solution with 50% methanol in water on each day of analysis. To determine their concentrations in mouse serum and tumor, their calibration standards were prepared by spiking 2 µL of KCI838 or KCI838 PME working solution into 38 µL of blank human plasma to make the final concentrations at 1, 2, 5, 10, 20, 50, 100, 200, 500, 1000, 2000 and 5000 nM. All standard samples were prepared freshly daily. 2.2.2. Sample processing for the determination of KCI838 and KCI838 PME serum and tumor concentrations

Frozen serum samples were thawed at room temperature, and an aliquot of 40 µL plasma was transferred into a micro-centrifuge tube. Blank pooled human plasma, which was used in preparation of standard, was also used to dilute plasma samples when necessary. An aliquot of 120 µL of ice-cold methanol was added into each tube to precipitate the protein content. The mixture was vortexed for 10 s and centrifuged at *21952 g* at 4 °C for 10 min. the supernatant was transferred into an autosampler vial and 5 µL was injected for LC-MS/MS analysis. Tumor tissue samples were thawed at ambient temperature. Tissue homogenate was prepared by addition of 4 volumes of PBS (*e.g.*, 400 µL PBS for 100 mg tissue) and homogenizing in a Precelly® homogenizer at *2000 g* for two 10 s sessions and 10 s pause. Tissue homogenate samples were further processed following the sample preparation method of plasma samples.

### Statistical Methods

For comparison of gene expression values, an unpaired t-test was used after log-transformation of expression measurements.

For comparison of paired experimental measurements in drug dose responses of promoter activity, BRET assay signal or hepatic enzyme induction, a paired t-test was used after log-transformation of response values.

For the MTT assays monitoring the time courses of cell growth inhibition using various cell lines and compounds at different doses, the drug treatment curves were compared in each case with the corresponding vehicle control using an unpaired t-test after log-transformation, where appropriate. The values of optical density were normalized across days by dividing by the mean value at Day 0.

IC50 and R^2^ values for colony growth inhibition and enzyme inhibition were calculated by fitting dose-response curves using nonlinear regression.

In all the *in vivo* drug efficacy studies in which mice were administered KCI838PME or placebo, the distributional assumption of tumor volumes was checked, and a square root transformation was applied when necessary to meet the normality assumption. Tumor volumes were measured in the left and right flanks of each mouse on days 0, 4, 6, and 8 (for bolus treatment) or on days 0, 7, 10, 12 and 14 (for controlled release treatment). Tumor growth curves were generated using the median and interquartile range interval. Tumor growth rates were compared using a linear mixed-effects model, with mouse and tumor side (left and right flanks) treated as random effects. Tumor volumes on days they were measured were compared using two-sample t-tests.

Where applicable, p-values were adjusted for multiplicity using Holm’s procedure. Statistical analyses were conducted using GraphPad Prism and the R statistical software packages.

## Results

### The novel KCI830 series of compounds

We synthesized a series of nine compounds with the core scaffold, 5-Hydroxy-2-(3-hydroxyphenyl)-1-methylquinolin-4(1H)-one in which the C ring nitrogen atom is attached to chemical groups with a range of hydrophobicity and steric features (**Figure 1A**) as described under Methods. They include substitution with a methyl group (KCI830), a propargyl group (KCI831), an ethyl group (KCI832), an n-propyl group (KCI833), an isopropyl group (KCI834), an isobutyl group (KCI835), a tert-butyl group (KCI836), a difluoro-ethyl group (KCI837) and a trifluoro-ethyl group (KCI838) (**Figure 1A**). The compound structures and their purity (>98%) were established, based on mass spectrometry, ^1^H NMR, ^13^C NMR, ^19^F NMR, HPLC and elemental (CHN) analysis.

**Figure 1:**
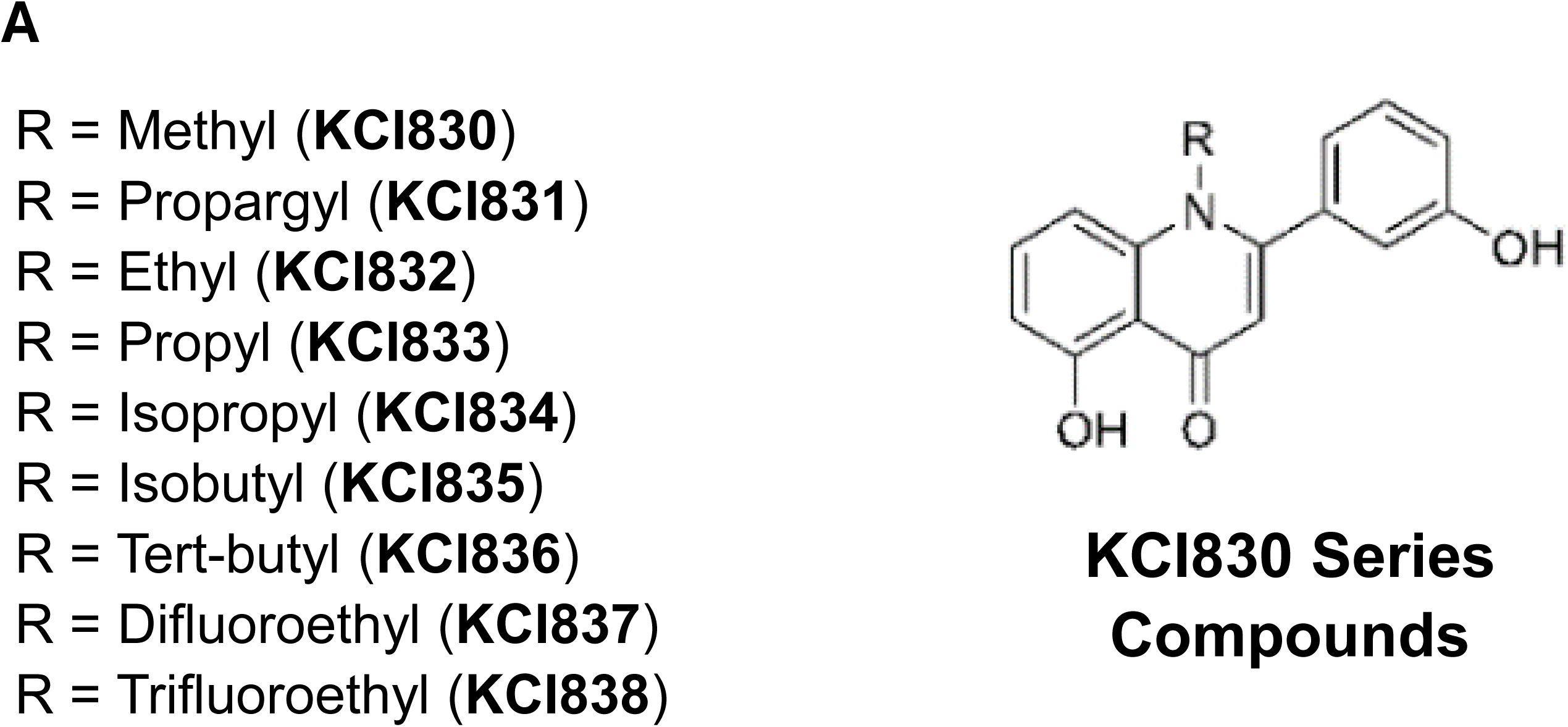
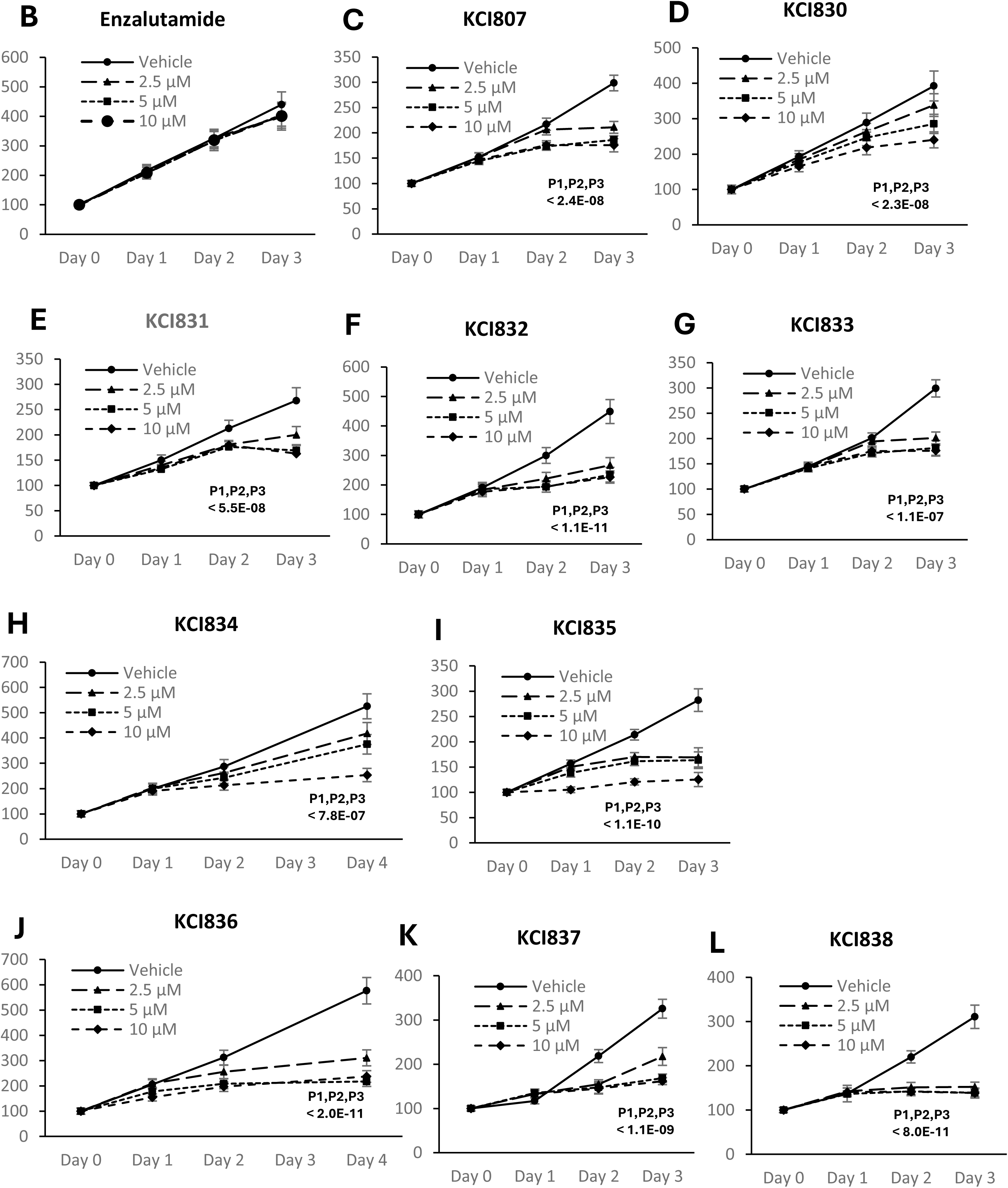
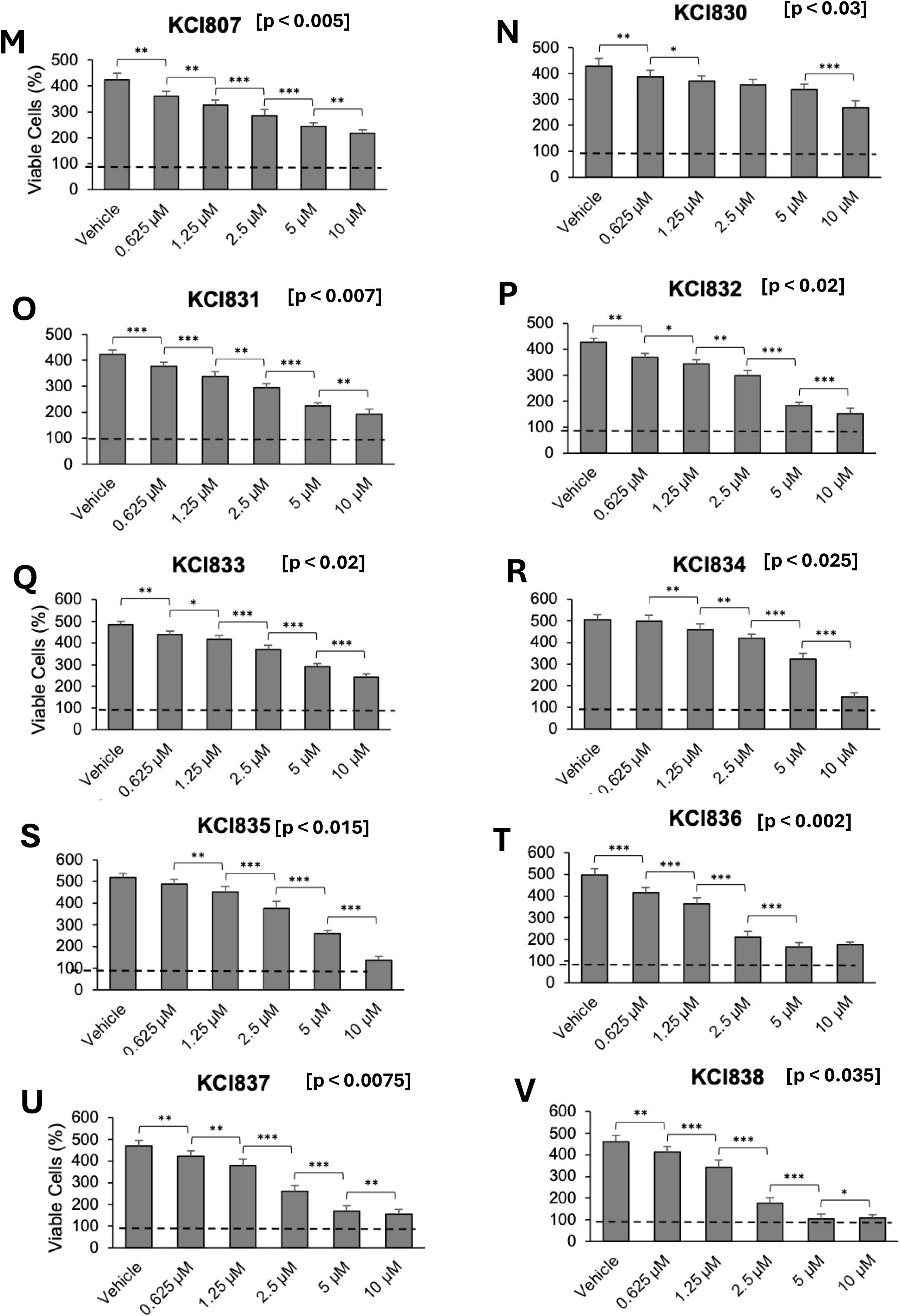
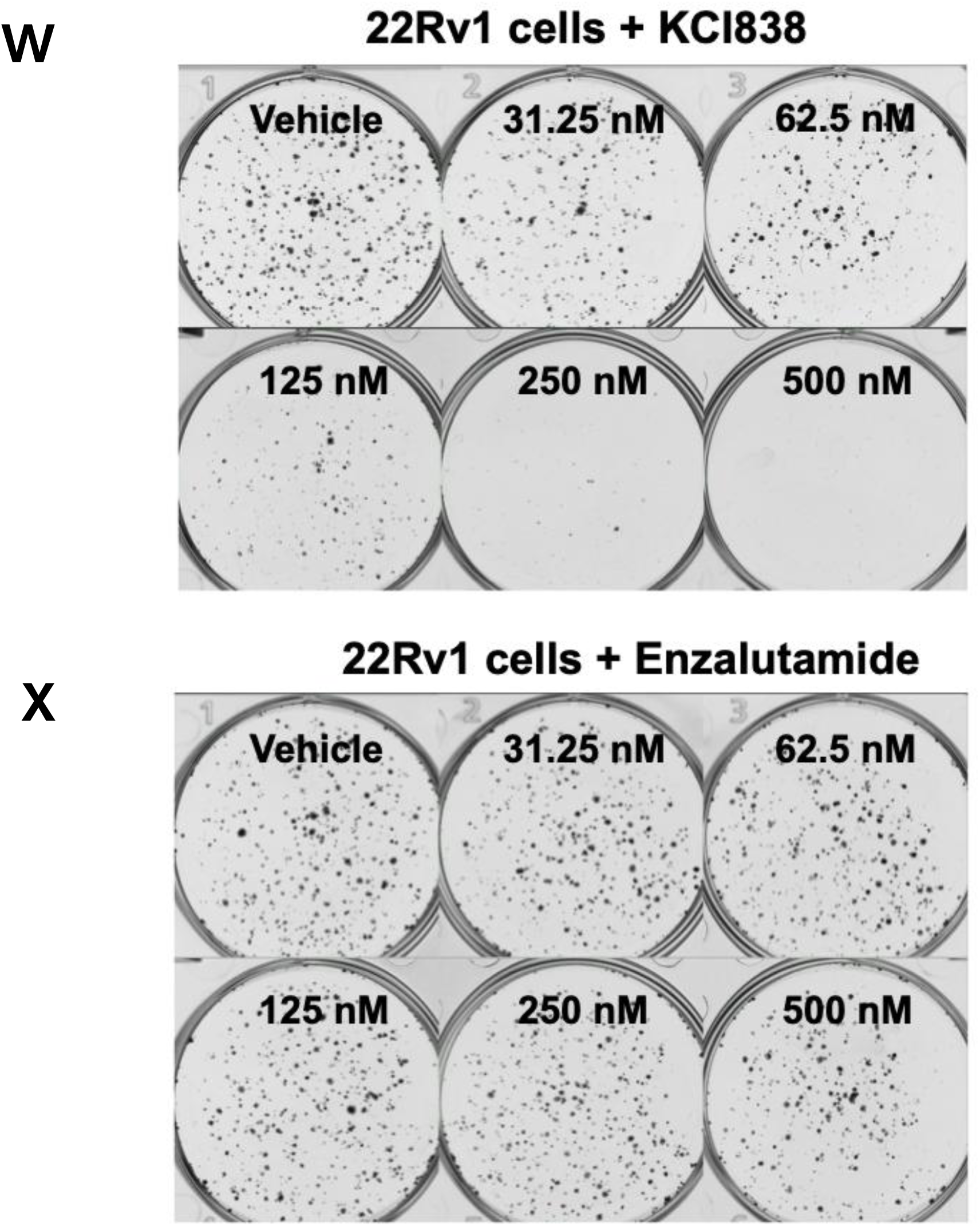
Structures of the KCI830 series of compounds and their ability to inhibit growth of 22Rv1 (CRPC) cells. ***Panel A:*** The core 5,3’-dihydroxy quinolone scaffold structure of the KCI830 series of compounds with the list of substituent groups on the C ring nitrogen atom that define members of the series. ***Panels B-L:*** Viability of 22Rv1 cells was determined at the indicated time points of treatment with vehicle or three concentrations of the various compounds, as indicated, using a standard MTT assay conducted as described under Methods and values were plotted as percent of the Day 0 values. For statistical analysis, the treatment curves were compared in each case with the corresponding vehicle control as described under Methods. ***Panels M-V:*** Viability of 22Rv1 cells was determined at the end of four days of treatment with vehicle and five concentrations of the various compounds, as indicated, using a standard MTT assay conducted as described under Methods and values were plotted as percent of the Day 0 values, considering the Day 0 values to be 100 percent. Statistical analysis of the histograms was conducted as described under Methods. ***Panel W, X:*** The effects of KCI838 or enzalutamide on colony formation by 22Rv1 cells in conditioned media, at the indicated concentrations of the compounds were tested as described under Methods.

### The KCI830 series of compounds have varying activities as inhibitors of the growth of castration resistant PCa (CRPC) cells

The compounds were screened for growth inhibitory activity in monolayers of the enzalutamide-resistant 22Rv1 CRPC cells, comparing the time courses of growth inhibition using 3 compound doses (2.5uM, 5 uM and 10uM) (**Figure 1B-L**). Whereas enzalutamide did not inhibit cell growth (**Figure 1B**), all the KCI830 series compounds showed inhibitory activity (**Figure 1D – 1L**), similar to KCI807 **(Figure 1C)**. KCI838 showed the earliest onset of growth inhibition among all the compounds, with virtually complete growth inhibition observed at all concentrations tested **(Figure 1L)**.

The dose-dependence of cumulative growth inhibition at the end of 4 days of treatment was compared for KCI807 **(Figure 1M)** and all the KCI830 series compounds **(Figure 1N – 1V)** in the dose range of 0.625uM - 10uM. KCI838 showed the most potent dose response for growth inhibition **(Figure 1V)**.

### KCI838 inhibits colony formation in enzalutamide resistant CRPC cells

Cell growth inhibitors often inhibit colony formation in a significantly lower dose range than growth inhibition in MTT assays as, in contrast to growth conditions in MTT assays, colony formation occurs over a longer period and involves many more replication cycles. In 22Rv1 CRPC cells, KCI 838 inhibited colony formation in the nanomolar dose range with the colony counts showing an IC50 of ∼60nM ( **Figure 1W**). In contrast, enzalutamide had no effect on colony formation in the same dose range (**Figure 1X**).

### KCI838 inhibits growth in other AR-dependent PCa cell line models better than KCI807 and enzalutamide

When tested in the dose range of 2.5uM to 10uM, KCI807, KCI838 and enzalutamide all inhibited monolayer growth of the androgen-dependent VCaP PCa cell line in a dose-dependent manner, with KCI838 showing better inhibition than either KCI807 or enzalutamide at the lower doses **(Figure 2 A-C)**. Dose-dependent inhibition of monolayer growth in the androgen-dependent LNCaP PCa cells was better for both KCI838 and KCI807, compared with enzalutamide (**Figure 2 D-F**). Additionally, in the CRISPR-generated CWR22Rv1-AR-EK PCa cell model that lacks full-length AR and is exclusively dependent on endogenous AR splice variants (AR-Vs), enzalutamide as expected was unable to inhibit growth, but the cells were sensitive to both KCI807 and KCI838, with KCI838 showing the stronger inhibition **(Figure 2 G-I)**.

**Figure 2:**
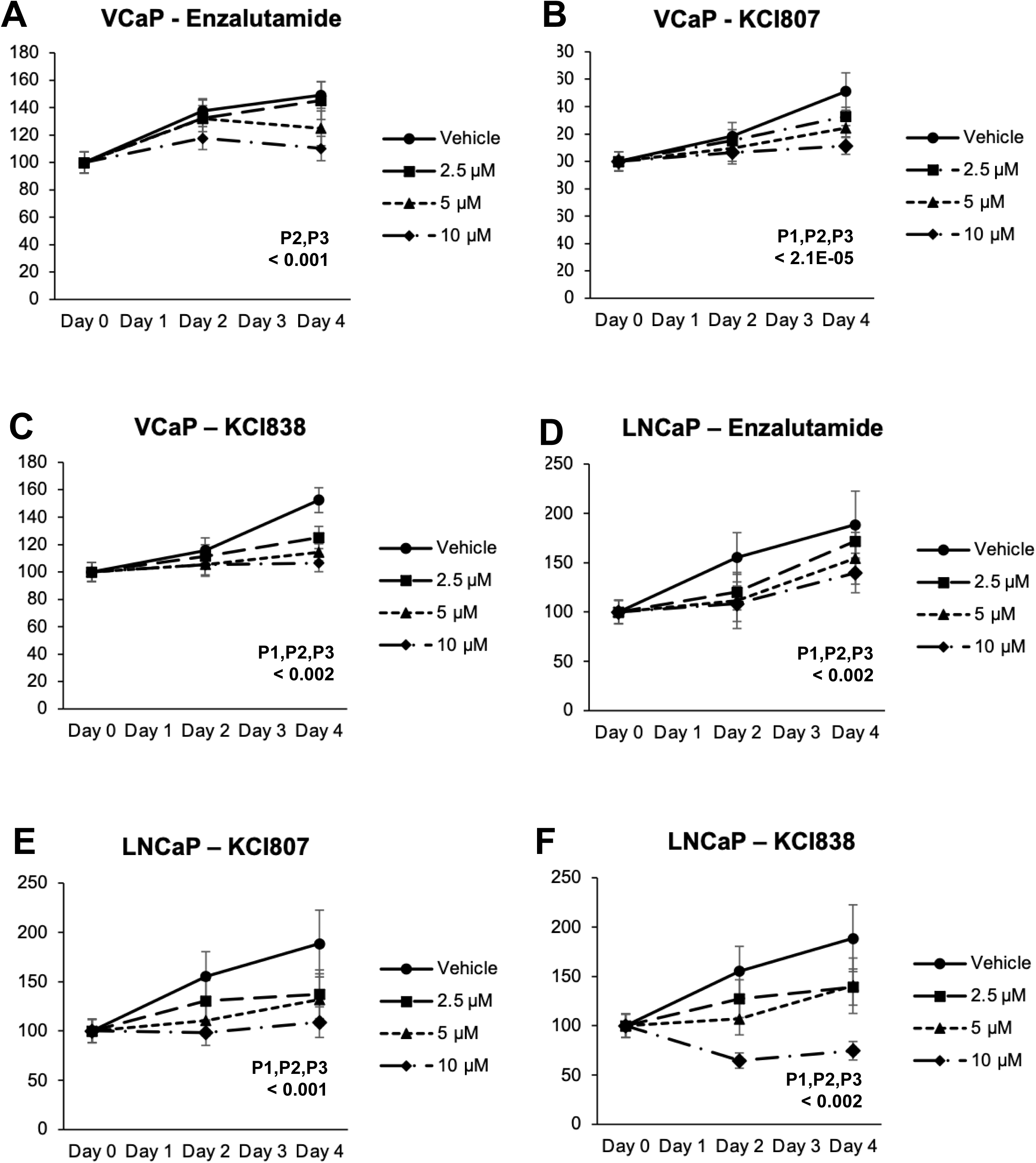
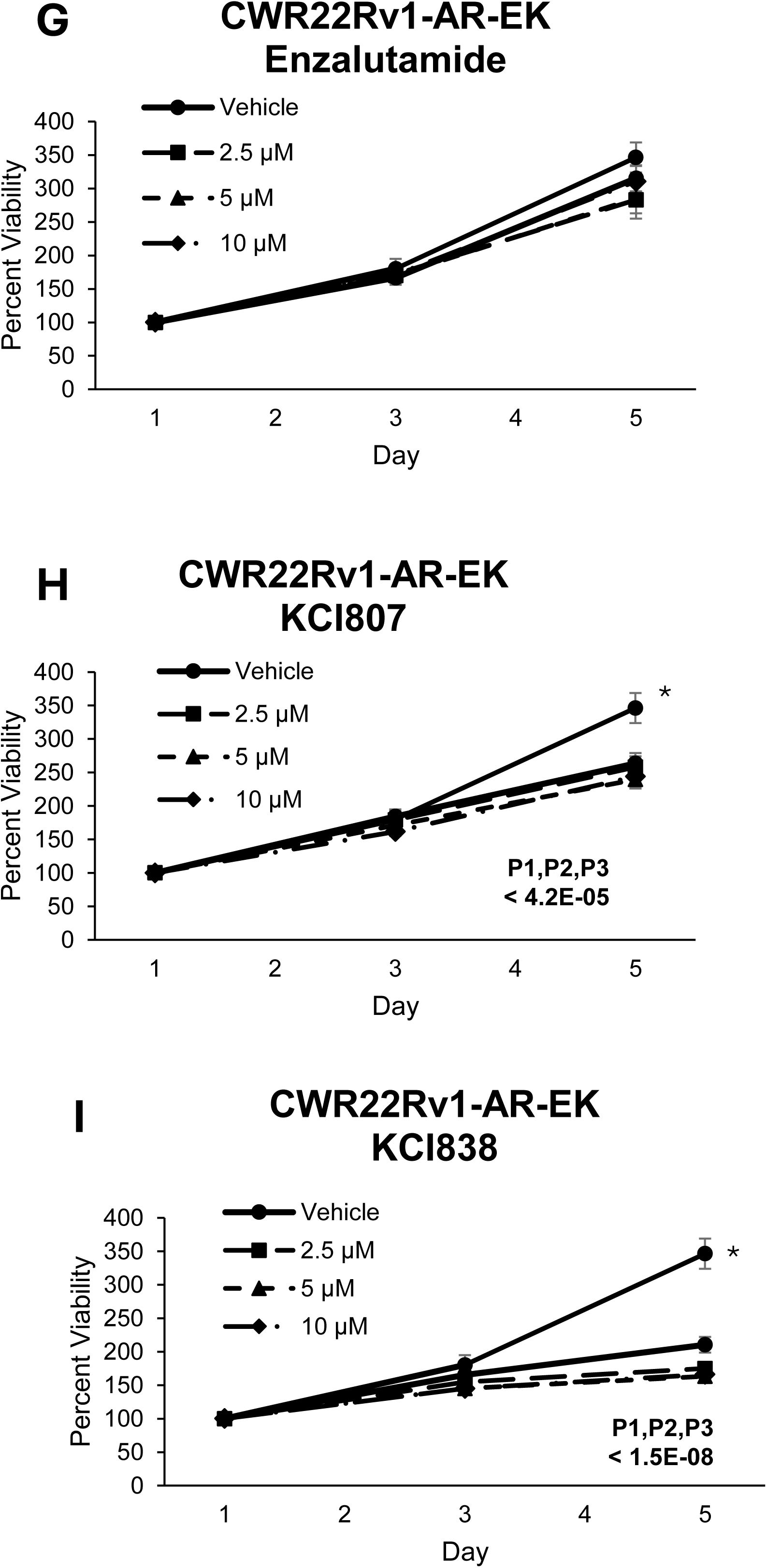
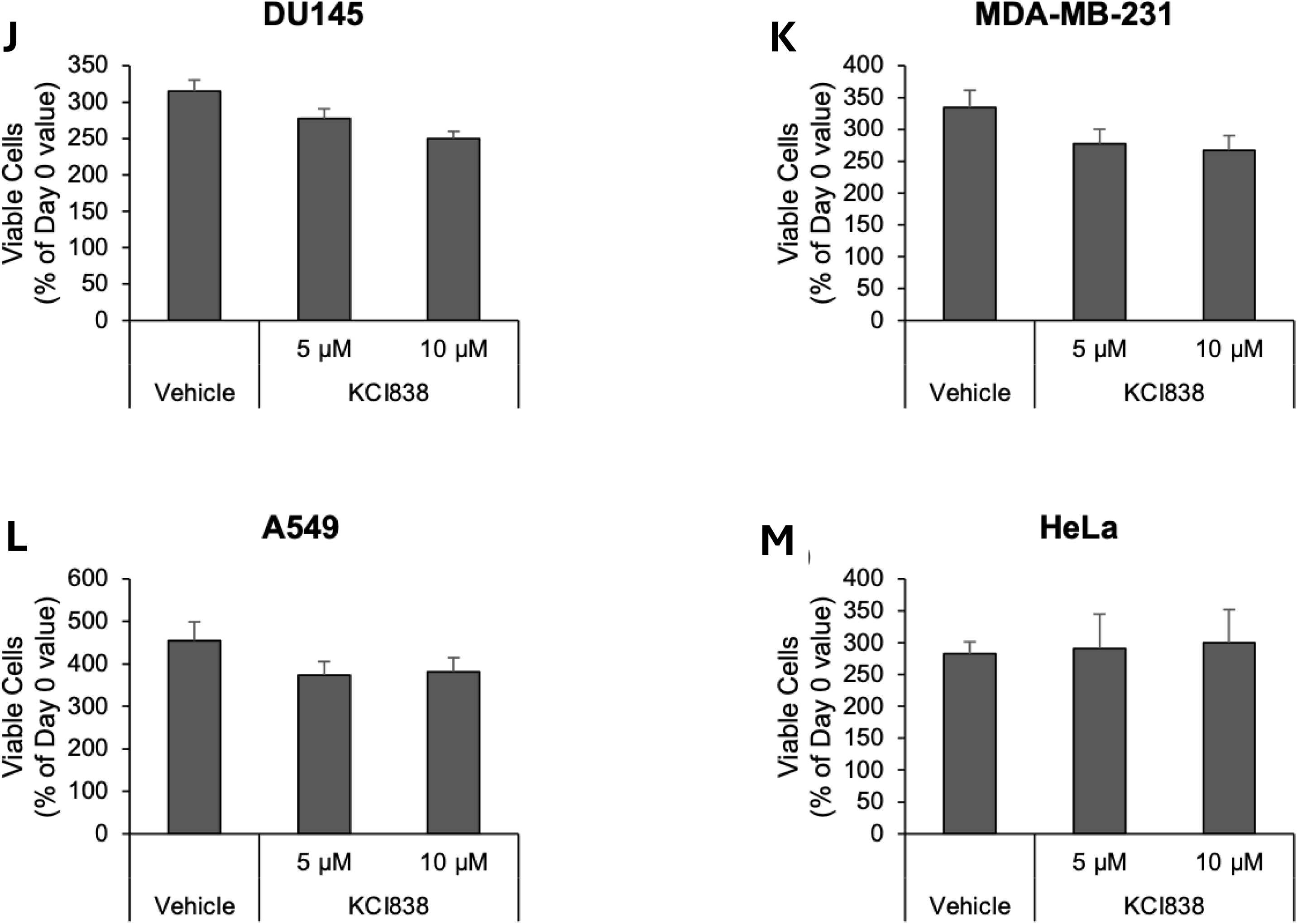
Selective growth inhibition by KCI838 of a variety of AR-positive prostate cancer cell line models but not AR-negative cells. ***Panels A-I:*** Viability of VCaP cells, LNCaP cells or CWR22Rv1-AR-EK cells was determined at the indicated time points of treatment with vehicle or three concentrations of the enzalutamide, KCI807 or KCI838, as indicated, using a standard MTT assay conducted as described under Methods and values were plotted as percent of the Day 0 values. For statistical analysis, the treatment curves were compared in each case with the corresponding vehicle control as described under Methods. ***Panels J-M:*** Viability of the AR-negative DU145 cells, MDA-MB-231 cells, HeLa cells and MDA-MB-231 cells was determined at the end of four days of treatment with vehicle and two concentrations of KCI838, as indicated, using a standard MTT assay conducted as described under Methods and values were plotted as percent of the Day 0 values, considering the Day 0 values to be 100 percent. Statistical analysis of the histograms, conducted as described under Methods, did not show significant effects of the compounds.

### KCI838 is inactive in AR-negative cell lines

The effect of KCI838 on monolayer growth was tested in four AR-negative cancer cell lines. KCI838 did not significantly affect growth of the AR-negative cell lines, including DU145 PCa cells (**Figure 2J**), MDA-MB-231 breast cancer cells (**Figure 2K**), A549 lung cancer cells (**Figure 2L**) and HeLa cervical cancer cells (**Figure 2M**).

### KCI838 selectively blocks the association of ELK1 and AR in situ

The ability of KCI838 to disrupt the binding of AR to ELK1 *in situ* was directly tested using a BRET assay. The BRET signal for binding of AR to ELK1 in situ was first measured as a function of the ratio of ELK1 to AR, ranging from a ratio of 0.5 to 12 (**Figure 3A, top**) and expression of the transfected proteins was confirmed by western blot (**Figure 3A, bottom**). Based on this data, an ELK1:AR ratio of 4 was used for testing the effect of KCI838 in **Figure 3B**. KCI838 strongly inhibited the BRET signal for binding of AR to ELK1 at the two doses tested (60% and 95% inhibition at 0.4 µM and 10 µM respectively, of KCI838) (**Figure 3B, top**). Protein expression in the transfected cells in Figure 3B was confirmed by Western blot (**Figure 3B, bottom**). To ensure that KCI838 was not toxic to the HEK293 cells, viability of the cells was tested following extended (3 days) treatment with 0.4 µM and 10 µM KCI838 and was found to be unaffected (**Figure 3C**).

**Figure 3:**
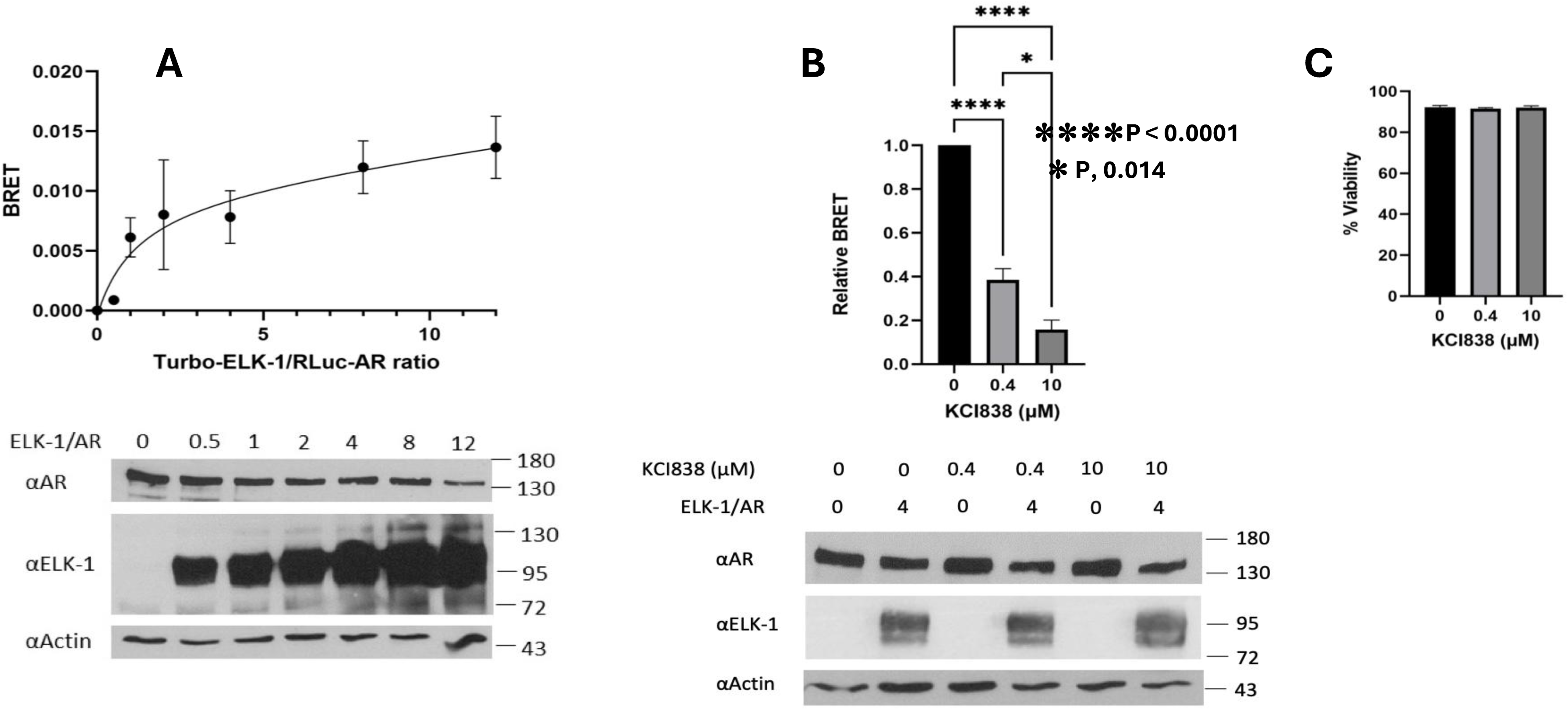
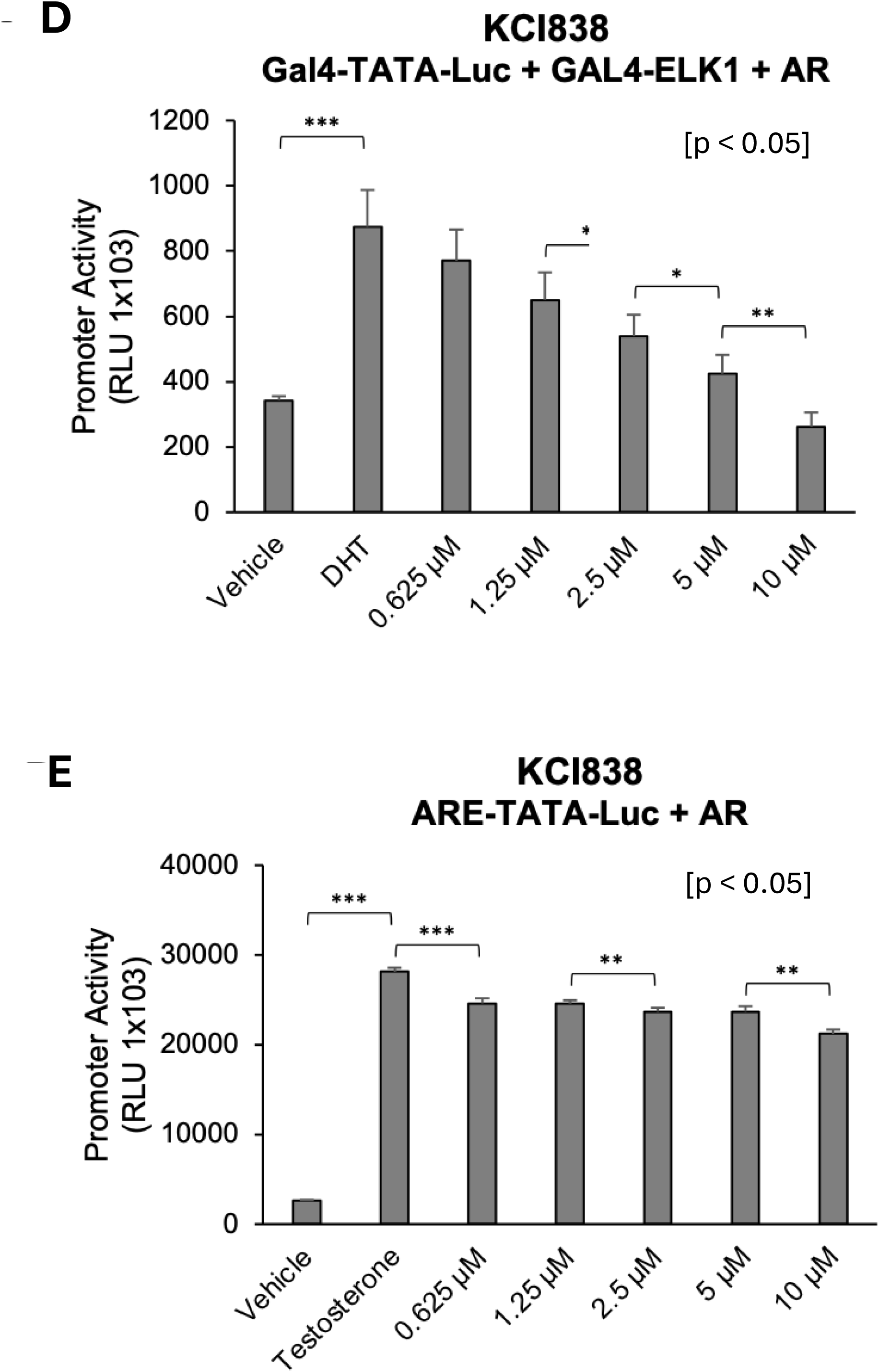
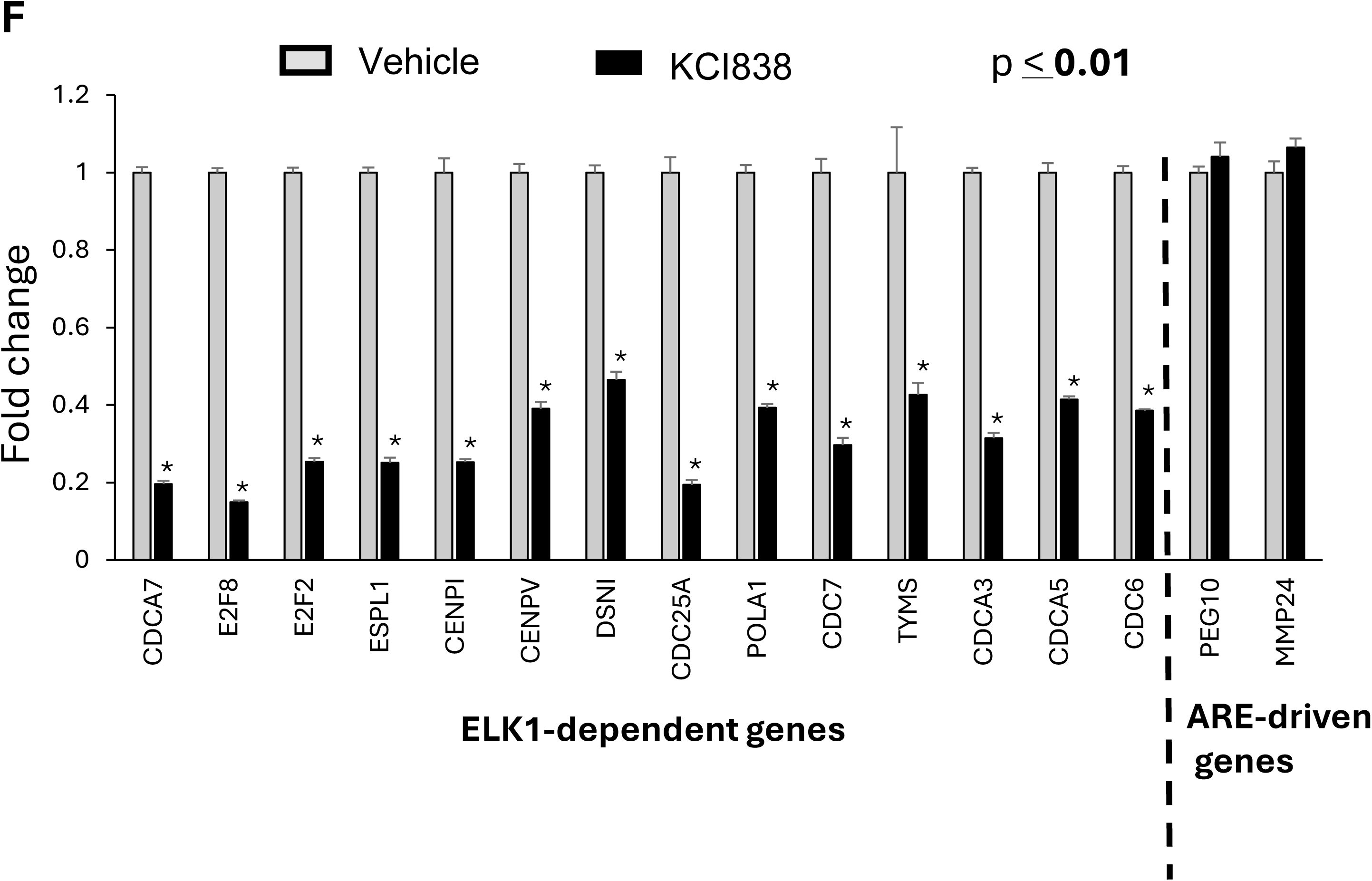
Blocking of ELK1 binding to AR *in situ* and preferential inhibition of ELK1-dependent promoter and gene activation by KCI838. ***Panel A:*** Optimization of the Turbo-ELK1:Rluc-AR ratio for the BRET assay to monitor in situ binding of ELK1 and AR in HEK293T cells was conducted as described under Methods. BRET readings were calculated as 635/535 ratio minus 635/535 ratio for Rluc-AR only transfected controls (Panel A, top). Cell lysates were probed by western blot for AR or ELK1, probing for β-actin as the loading control (Panel A, bottom). An ELK1:AR ratio of 4:1 was determined to be the optimal ratio for the inhibition assay in Panel B. ***Panel B:*** BRET assay was conducted as described under Methods, using a fixed ELK1:AR ratio of 4:1. 24 hours after transfection, cells were treated with DMSO (vehicle control) or KCI838 (0.4uM and 10uM) and left for 24 hours. They were then trypsinized and seeded in 96 well white-bottomed plates at 100,000 per well (in quadruplicate) in media lacking phenol red (with DMSO or drug as appropriate). After 2h, the BRET assay was completed as described under Methods (Panel B, top). Cell lysates were probed by western blot for AR or ELK1, probing for β-actin as the loading control (Panel B, bottom). Statistical analysis was conducted as described under Methods. ***Panel C:*** Cell viability in the presence of KCI838 was ensured by treating HEK293T cells (3×10^5^ cells/ml seeded in 6-well dishes 24 hours pre-treatment) for an extended period (3 days) with the compound or vehicle control and measuring viability using trypan blue staining. ***Panel D:*** The dose-dependent effect of KCI838 on ELK1-dependent promoter activation by androgen-activated AR was measured as described under Methods, using recombinant HeLa cells which have a stably integrated minimal promoter-luciferase reporter containing five upstream Gal4 elements (Gal4-TATA-Luc) and constitutively express a GAL4-ELK1 fusion protein (in which the Gal4 DNA binding domain is substituted for the ETS DNA binding domain of ELK1), as well as full-length AR. Statistical analysis was conducted as described under Methods. ***Panel E:*** The effect of a range of concentrations of KCI838 on ARE-dependent promoter activation by androgen plus AR was examined as described under Methods using recombinant HeLa cells harboring a minimal promoter-luciferase reporter with an upstream ARE sequence (ARE-TATA-Luc) and also stably expressing full-length AR. Statistical analysis was conducted as described under Methods. ***Panel F:*** The ability of KCI838 to inhibit ELK1-dependent vs. ARE-dependent endogenous AR target genes was examined in 22Rv1 cells, as described under Methods. The target gene sets chosen for this experiment are representative genes from the entire sets of ELK1-dependent or ARE-dependent androgen target genes that have been previously established. Statistical analysis was conducted as described under Methods.

### ELK1-dependent transcriptional activation by AR is inhibited by KCI838

The ability of KCI838 to selectively disrupt functional interactions between AR and ELK1 is illustrated in **Figure 3D-3F**. Recombinant HeLa cells harboring a Gal4-containing minimal promoter-luciferase reporter construct (Gal4-TATA-Luc) and stably expressing ectopic GAL4-ELK1 fusion protein as well as ectopic AR were used in **Figure 3D**. In these cells, promoter activation by androgen was inhibited by KCI838 in a dose-dependent manner (**Figure 3D**). In contrast, in recombinant HeLa cells harboring a classical androgen response element (ARE)-driven promoter (ARE-TATA-Luc) and stably expressing ectopic AR, KCI838 at best only minimally inhibited promoter activation by androgen (**Figure 3E**). Further, when 22Rv1 prostate cancer cells were treated with KCI838, the compound inhibited the expression of mRNAs of representative genes from the previously established set of ELK1-dependent endogenous target genes of AR (**Figure 3F**). In contrast expression of known classical ARE-driven gene targets of AR that are not associated with ETS elements or other possible AR tethering sites was unaffected by KCI838 (**Figure 3F**).

### The growth inhibitory effect of KCI838 is exclusively mediated by AR

To rule out the possibility of any off-target effects of KCI838 as mediating its growth inhibitory activity, we generated recombinant 22Rv1 cells in which total cellular AR was increased by lentiviral transduction and antibiotic selection of the pool of transduced cells as well as control parental cells transduced with the vector alone, followed by similar selection. The AR transduced cells (22Rv1-Lenti AR) showed a 2.1-fold higher level of total AR as measured by Western blot compared with the control vector transduced cells (22Rv1-Lenti vector) (**Figure 4A**). If AR is the mediator of the growth inhibitory effect of KCI838, then it would be predicted that for a 2-fold increase in AR expression, there should be no inhibition of growth in the AR overexpressing cells at a KCI838 concentration corresponding to the IC50 value for the control cells; this is because the total free AR at 50 percent occupancy by inhibitor in the 2x AR overexpressing cells should equal the total free AR in the untreated control cells. Additionally, the IC50 value of the compound in the AR overexpressing cells should correspond to the IC75 value for the control cells. To test this, the KCI838 dose responses for inhibition of colony formation were compared between the control ( **Figure 4B**) and AR overexpressing (**Figure 4C**) cells, treating with DHT in both cases to ensure efficient nuclear localization of the ectopic AR. As seen in **Figure 4B**, the IC50 and IC75 values were 68 nM and 121 nM, respectively, for the control cells. In the case of the AR overexpressing cells, there was no inhibition at 68 nM and the IC50 value increased to 132 nM, close to the IC75 value observed in the control cells (**Figure 4C**). These results clearly implicate AR as the principal or only functionally relevant target for the observed physiological activity of KCI838 in PCa cells.

**Figure 4:**
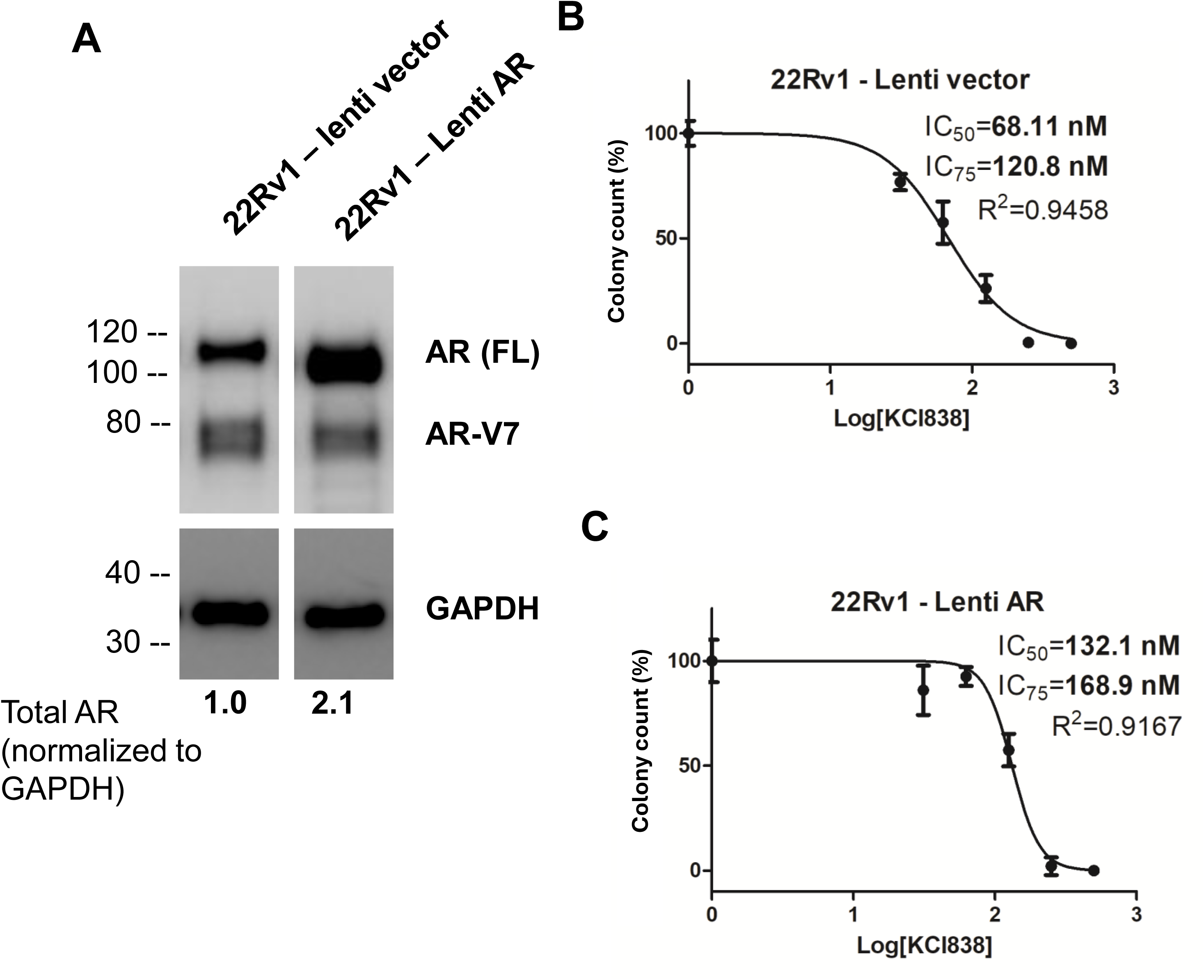
Modulation of physiological sensitivity to KCI838 by ectopic AR. ***Panel A:*** Western blot of cell lysates from recombinant 22Rv1 cells generated as described under Methods, by stable transduction with a lentivirus vector expressing full-length AR (22Rv1-Lenti AR) or the lentivirus vector alone (22Rv1-Lenti vector). The blots were also probed with antibody to GAPDH. The band intensities were quantified using Image Lab from Bio-Rad. The relative band intensity for total AR in the 22Rv1-Lenti AR cells vs. the 22Rv1-Lenti vector cells is indicated, after normalizing to the intensities of the GAPDH bands. ***Panel B-C:*** Colony formation of 22Rv1-Lenti vector cells or 22Rv1-Lenti AR cells was measured as described under Methods, after treatment of the plated cells with various concentrations of KCI838. The test compounds and media were replenished every 48 hours until colonies developed in the control untreated wells (in 7 - 10 days). IC50, IC75 and R2 values for colony growth inhibition were calculated by fitting dose-response curves using nonlinear regression.

### KCI830 series compounds are poor inducers and substrates for major human hepatic enzymes that metabolize drug molecules

**Figure 5** shows induction of major hepatic enzymes by KCI807 and KCI838 when primary human hepatocytes were exposed to the compound for 72 hours. Among monooxygenases, KCI807 strongly induced CYP1A2 at a dose of 0.5 µM, with maximum induction at 5 µM (**Figure 5A**). In contrast, KCI838 showed no induction at a dose of 0.5 µM and relatively low inhibition at 5 µM. Likewise, compared with KCI807, KCI838 was a relatively poor inducer of the monooxygenase CYP2B6 **(Figure 5B)**. Other monooxygenases, including CYP2A6, CYP3A4, CYP2C9 and CYP2C19 were not induced by either compound. The major UDP-glucuronyltransferases (that glucuronylate drugs) UGT1A1 **(Figure 5C)** and UGT1A4 **(Figure 5D)** were not induced significantly by KCI838 at doses of 0.5 µM and 5 µM in contrast to KCI807, which strongly induced these enzymes. Other UDP-glucuronyltransferases, including UGT1A6, UGT1A9 and UGT2B4 were not significantly induced by either compound. Induction of sulfotransferases (SULT1A1, SULT2A1) or glycosyltransferase (UGT2B7) by either compound was insignificant.

**Figure 5:**
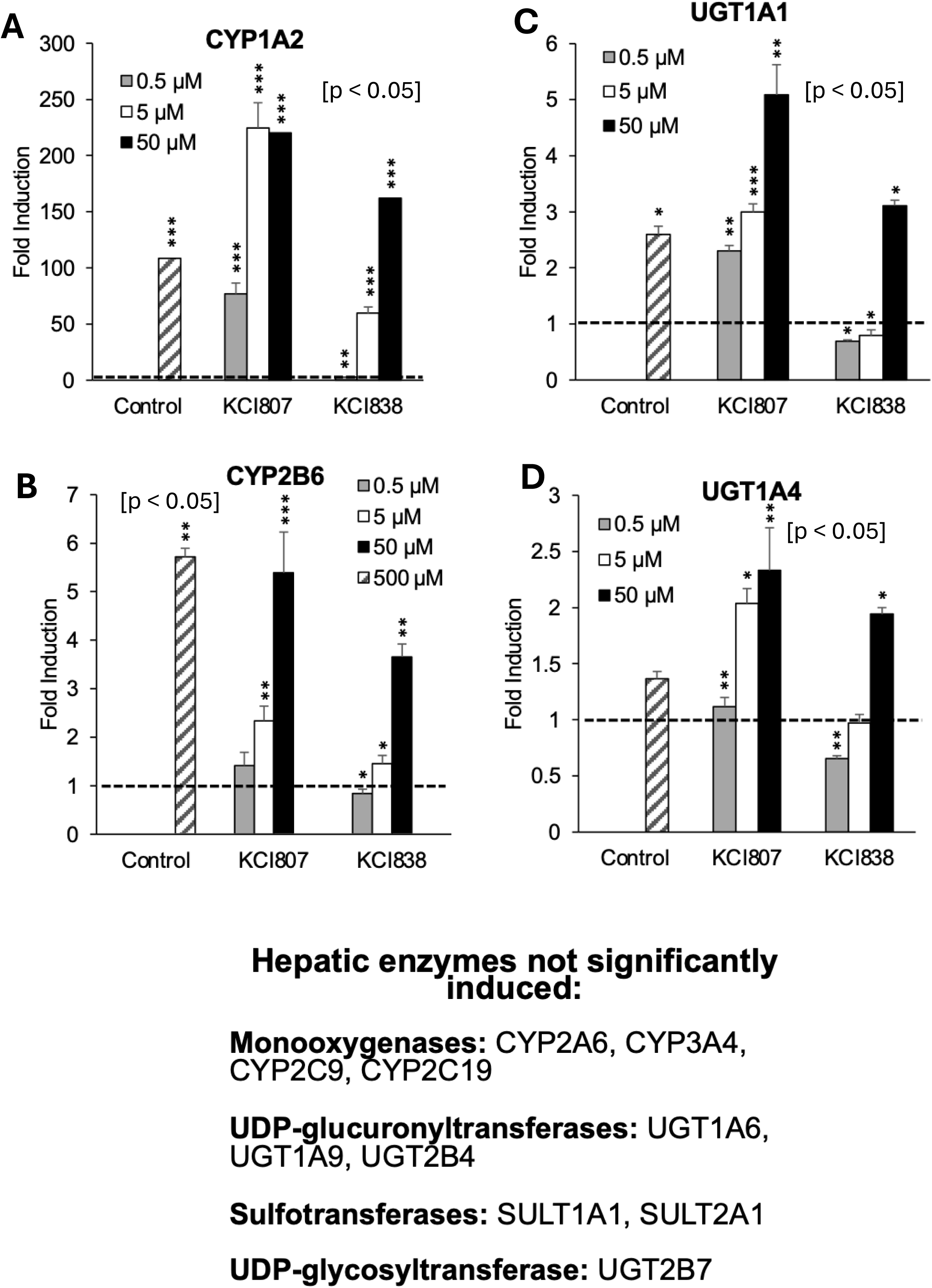
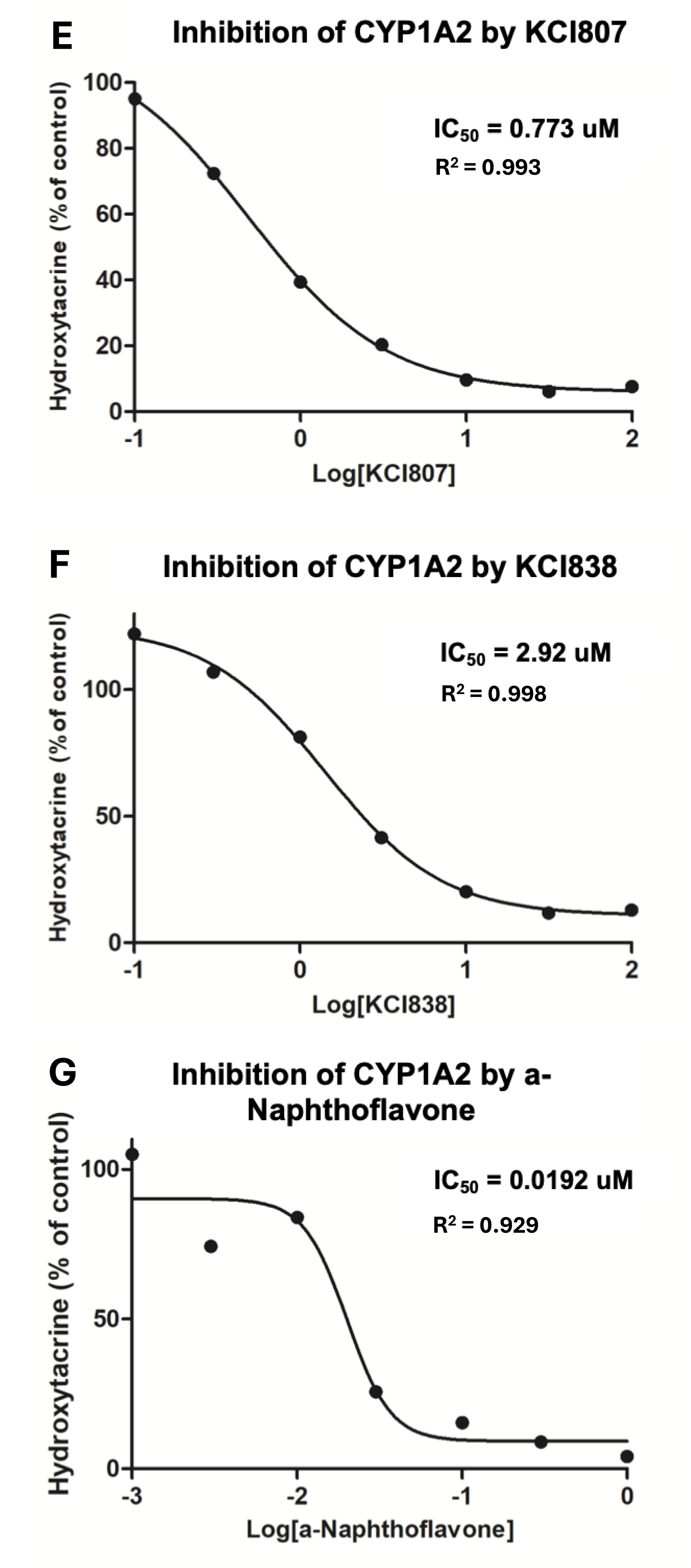
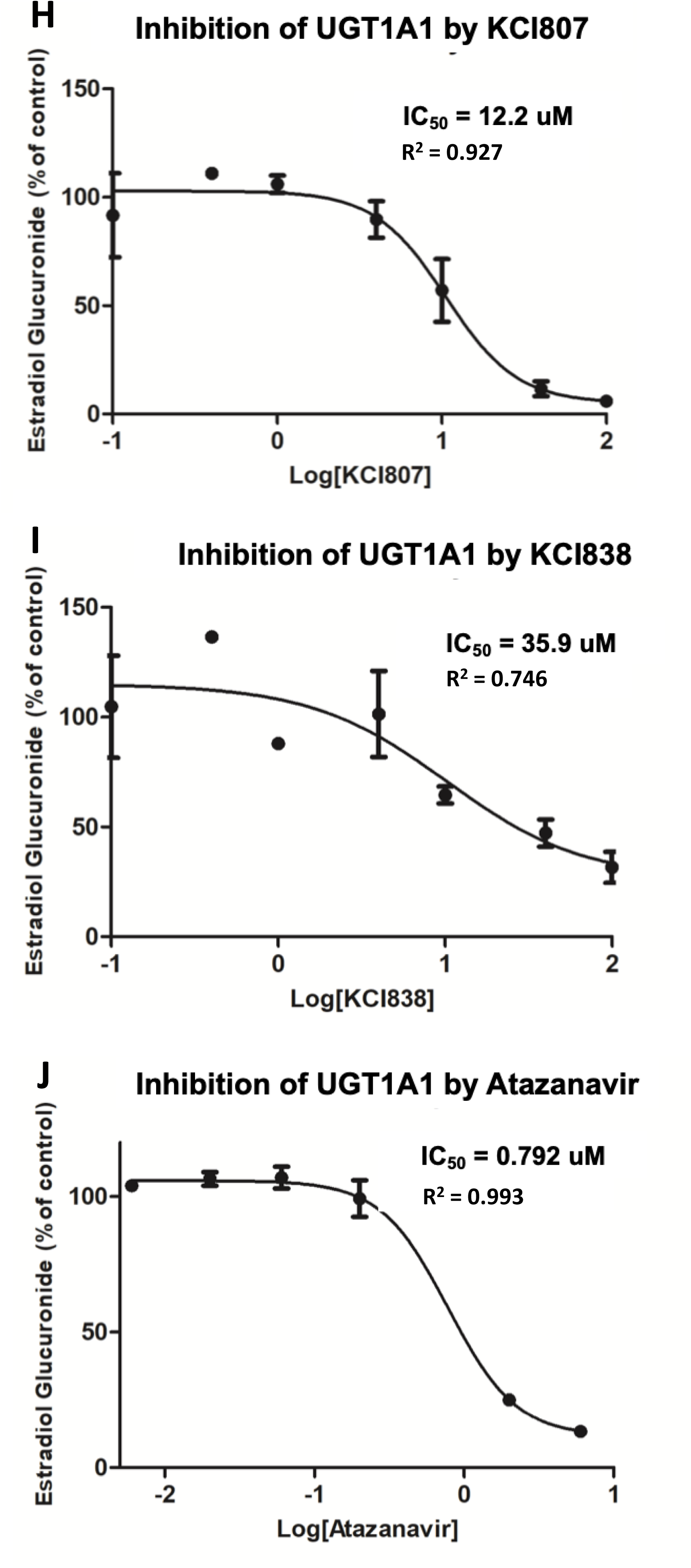
The relative ability of KCI838 to induce hepatic enzymes and to interact with CYP1A2 and UGT1A1. ***Panels A-D:*** Enzyme induction in primary human hepatocytes following 72h of treatment with positive control inducers (b-naphthoflavone, phenobarbital, dexamethasone and rifampin), KCI807 (0.5uM, 5uM and 50uM) or KCI838 (0.5uM, 5uM and 50uM) was measured as described under Methods. The mRNAs for the following hepatic enzymes were quantified: CYP1A2, CYP2A6, CYP2B6, CYP2C9, CYP2C19, CYP3A4; Sulfotransferases: SULT1A1, SULT2A1; UDP-glucuronyltransferases: UGT1A1, UGT1A4, UGT1A6, UGT1A9, UGT2B4; UDP-glycosyltransferase: UGT2B7. Statistical analysis was performed as described under Methods. ***Panels E-G:*** An established assay was used to compare inhibition of CYP1A2 by KCI838 vs. KCI807. This assay was conducted by Cyprotex US LLC (Watertown, MA) as described under Methods. The selective CYP1A2 inhibitor α-naphthoflavone was used as a positive control. Inhibition was monitored as the decrease in the formation of the metabolite compared to vehicle control. IC50 and R2 values for enzyme inhibition were calculated by fitting dose-response curves using nonlinear regression. ***Panels H-J:*** An established assay was used to compare inhibition of UGT1A1 by KCI838 vs. KCI807. This assay was conducted by Cyprotex US LLC (Watertown, MA) as described under Methods. The UGT1A1 inhibitor, atazanavir, was used as a positive control. The decrease in the formation of the metabolite compared to the vehicle control was used to monitor enzyme inhibition. IC50 and R2 values for enzyme inhibition were calculated by fitting dose-response curves using nonlinear regression.

### KCI838 interacts poorly with CYP1A2 and UGT1A1 compared with KCI807

The interactions of KCI838 and KCI807 with the major human liver monooxygenase CYP1A2 (**Figure 5E-5G**) and the major human UDP-glucuronyltransferase UGT1A1 (**Figure 5H-5J**) (i.e. the ability of the compounds to serve as inhibitor or substrate for the enzymes) was by determining the IC50 values of the compounds in standard enzymatic activity assays using the specific substrates noted in the Figure legends. The IC50 value of KCI838 was relatively high for CYP1A2 (3 µM) **(Figure 5F)** compared with KCI807 (0.8 µM) **(Figure 5E)** and the control inhibitor (0.02 µM) **(Figure 5G)**. The IC50 value of KCI838 was also relatively high for UGT1A1 (36 uM) **(Figure 5I)** compared with KCI807 (12 µM) **(Figure 5H)** and the control inhibitor (0.8 µM) **(Figure 5J)**.

### KCI838 effectively and reversibly inhibits growth of aggressive enzalutamide-resistant patient-derived tumor xenografts

To test its antitumor activity *in vivo*, KCI838 was converted to a soluble prodrug form. Accordingly, KCI838 was synthesized as the disodium salt of a phosphate monoester at the 3’ hydroxyl position (KCI838PME) (Figure 6A). The rationale for conversion to the phosphate ester is that 1) The hydrophobicity of KCI838 limits bioavailability when administered systemically, because of its relatively poor solubility and 2) Primary and metastatic human PCa tumors are characteristically rich in a secreted acid phosphatase (prostatic acid phosphatase) that should hydrolyze the prodrug, releasing the active KCI838 at the tumor site that should then be efficiently absorbed by the tumor tissue due to its hydrophobicity. A patient-derived tumor xenograft (PDX) model had to be used for this purpose, as in contrast to clinical tumors, established PCa cell lines express low or absent levels of phosphatases. The PDX model (PDX-PR011) has been reported previously as an aggressive tumor that is AR-positive and enzalutamide-resistant (27). The tumor was also rich in prostatic acid phosphatase, as determined by immunohistochemistry (Figure 6B). As the solubility of the prodrug should limit systemic retention time due to rapid renal clearance, we postulated that the anti-tumor efficacy of KCI838PME would be optimal when administered using a controlled release method vs. daily bolus injections.

**Figure 6:**
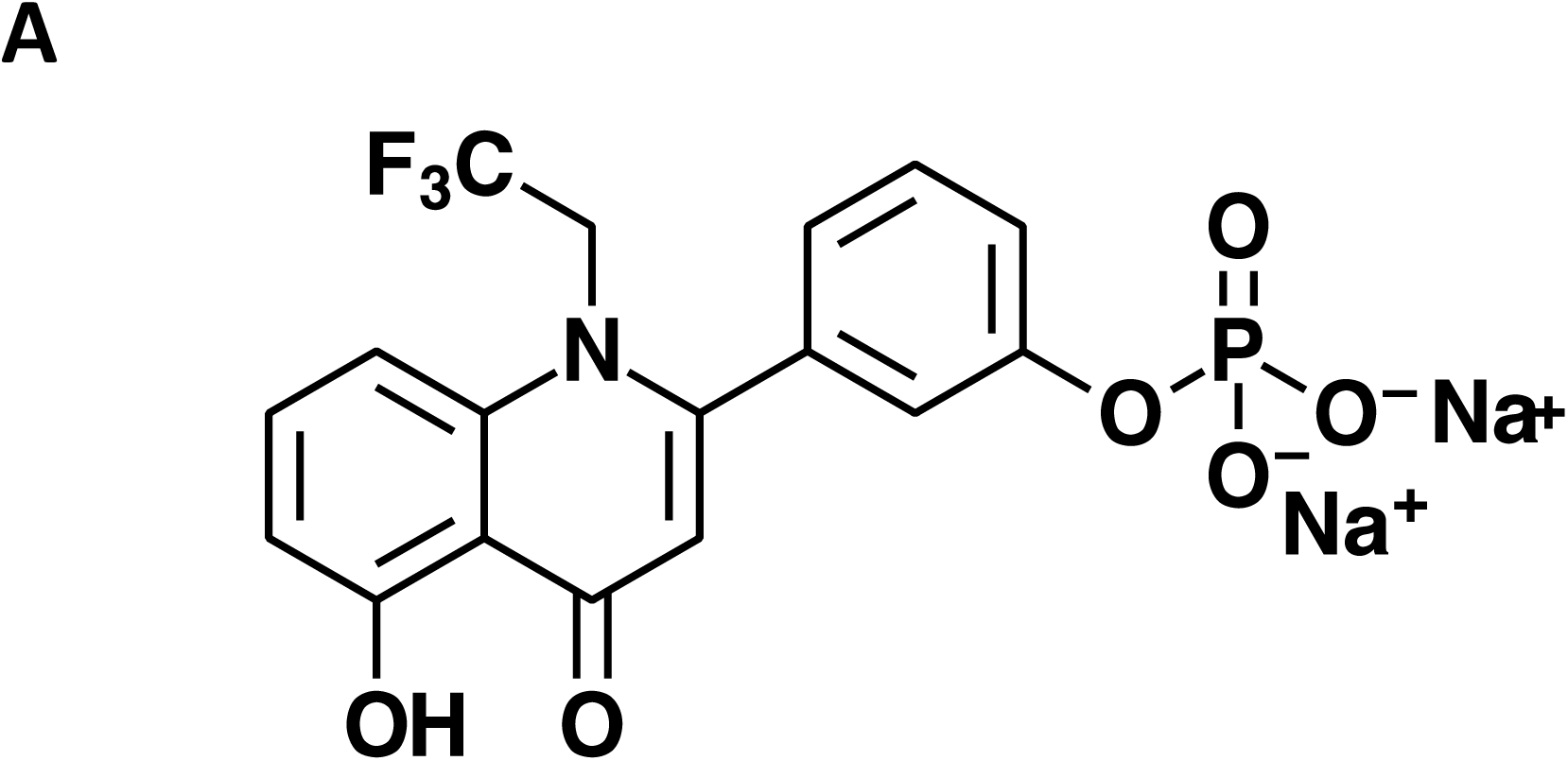
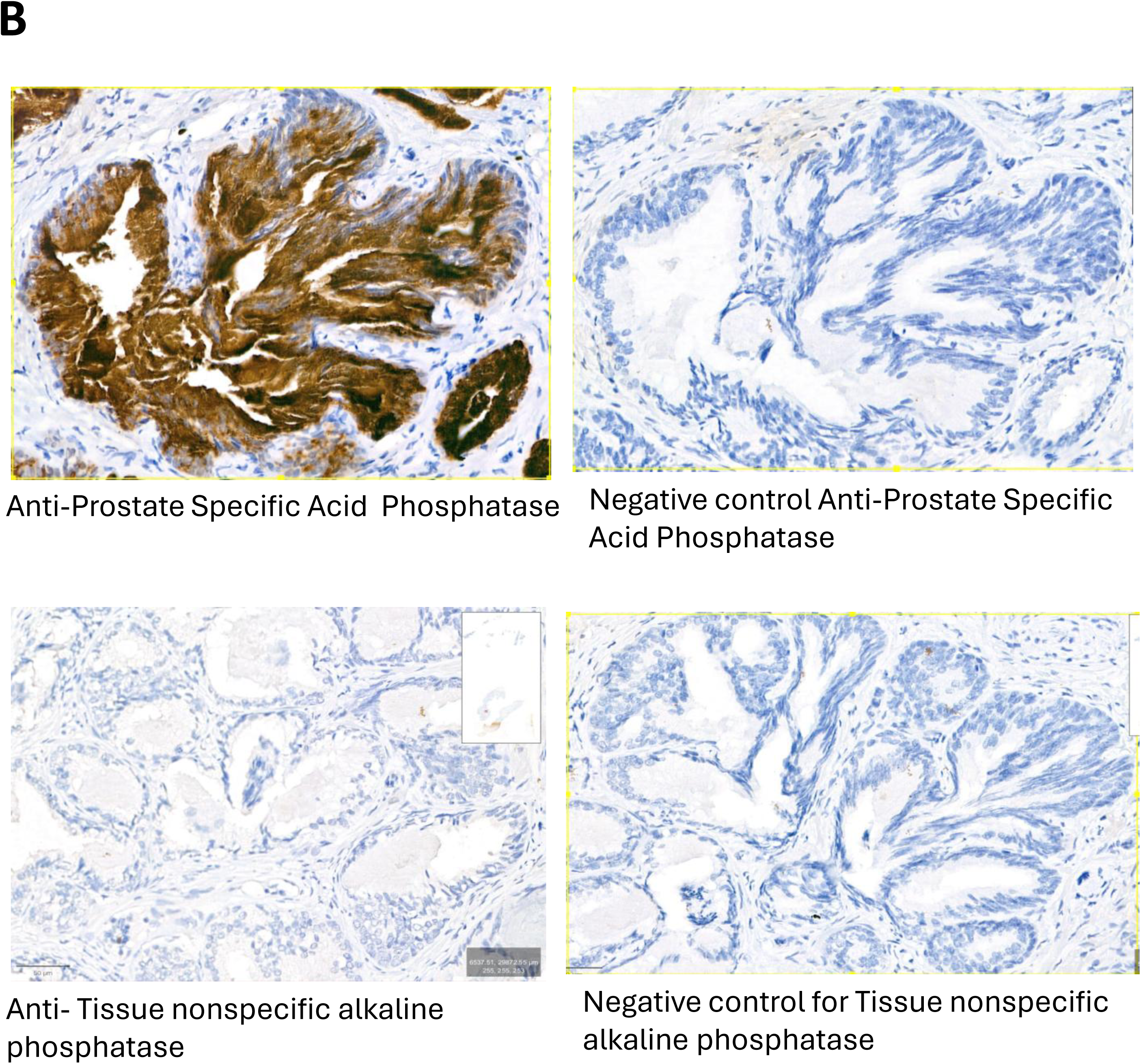
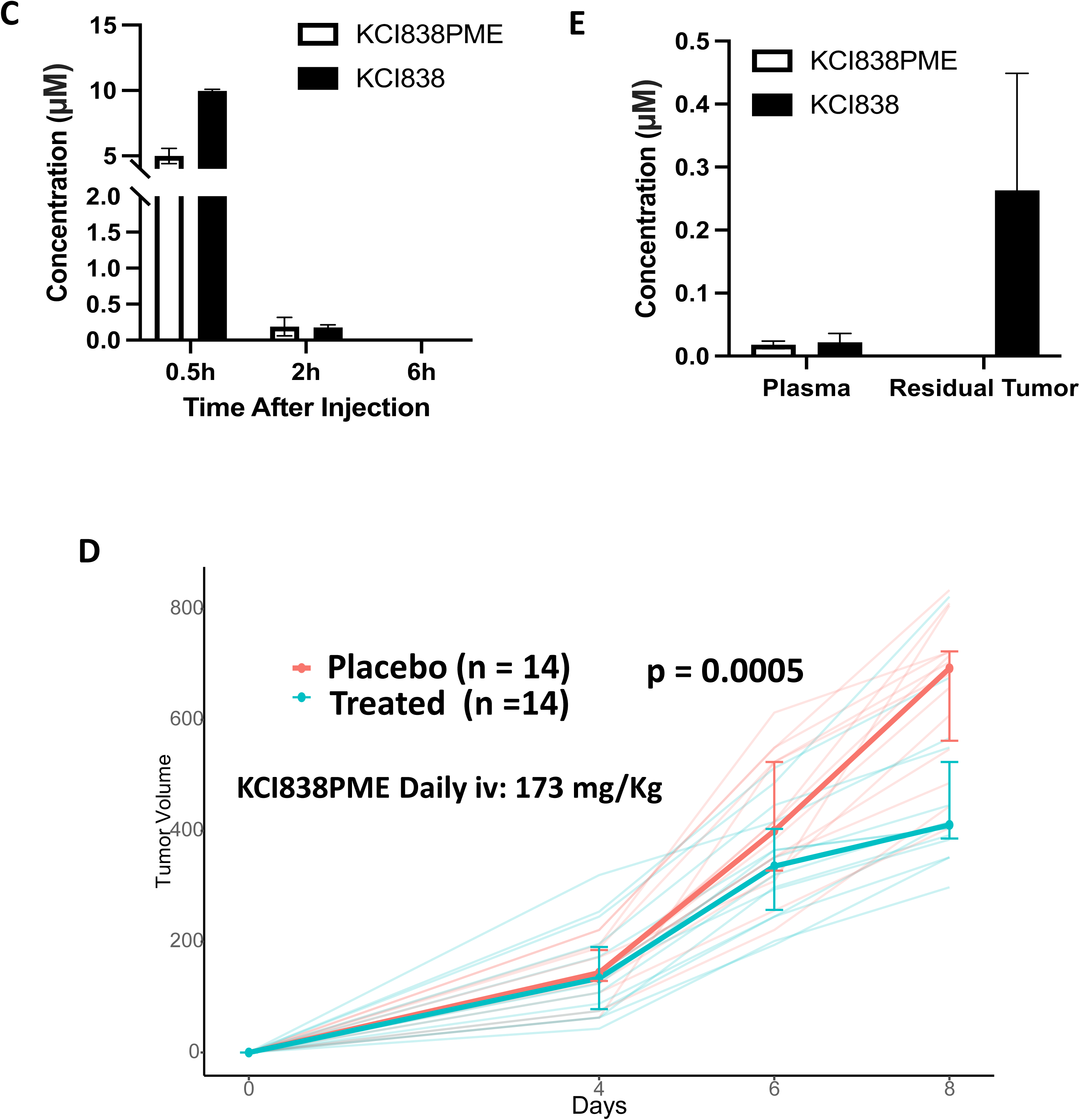
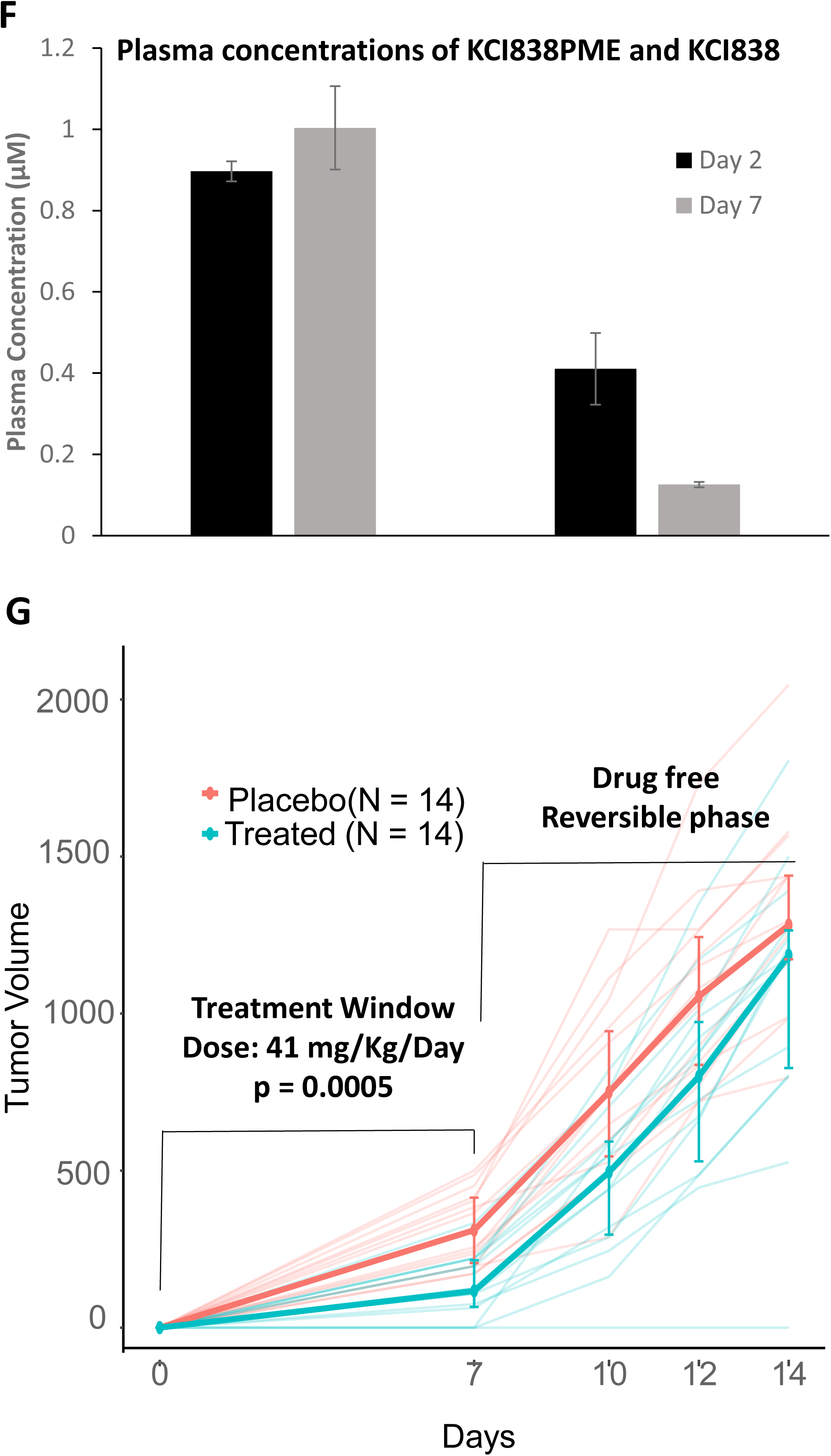
Tumor growth inhibition by KCI838PME in a prostate cancer PDX model. ***Panel A:*** Structure of the KCI838 3’-monophosphate ester (KCI838PME) prodrug. ***Panel B:*** Expression of Prostatic Acid Phosphatase in the PDX-PR011 tumor. Immunohistochemistry staining of paraffin embedded sections of the human PDX tumor tissue was conducted by iHisto Inc. as described under Methods. ***Panel C:*** Nine non-tumor bearing mice were injected 180 mg/kg of KCI838PME iv in water and blood samples drawn from groups of 3 mice at 0.5h, 2h and 6h for preparation of plasma. The plasma samples were analyzed by LC/MS to measure circulating levels of KCI838PME and KCI838, as described under Methods. ***Panel D:*** The PDX-PR011 tumor was implanted sc, bilaterally in groups of 7 mice. The treated group was injected daily on Days 1-8 with KCI838PME. The bolus injections were administered iv at a drug dose of 4mg per mouse per day in water. Treatment with KCI838PME was initiated within 1 day of tumor implantation. Tumor measurements and endpoint determination are described under Methods. ***Panel E:*** Six hours after the 8^th^ bolus injection, in Panel D, the mice were sacrificed after drawing blood samples for preparation of plasma and measurement of plasma levels of KCI838PME and KCI838 by LC/MS as described under Methods. At the same time, the residual tumors were harvested to measure tumor levels of KCI838PME and KCI838 by LC/MS as described under Methods. ***Panel F:*** Groups of 3 non-tumor bearing mice were implanted primed ALZET pumps (2 pumps per mouse, containing a total of 8 mg KCI838PME in water, per mouse). Two days and 7 days after implantation of the pumps, plasma was obtained from the mice to measure levels of KCI838PME and KCI838 by LC/MS as described under Methods. ***Panel G:*** The PDX-PR011 tumor was implanted sc, bilaterally in groups of 7 mice on Day 0. On Day 1, the mice were implanted with two primed ALZET pumps per mouse (1-week release, 200 ul capacity) containing either water (control group) or a 20mg/ml solution of KCI838PME in water (total, 8mg KCI838PME per mouse in the treatment group). The drug was exhausted on Day 8 but all mice were kept until Day 14. Tumor volumes were measured as described under Methods. In ***Panel D*** and ***Panel G***, statistical analyses of the tumor growth curves were performed as described under Methods.

First, using non-tumor bearing mice, it was determined that following a single bolus IV injection of 180 mg/kg of KCI838PME, the average plasma levels of KCI838PME and KCI838 at 0.5 hours, 2 hours and 6 hours were, 5.2 µM, 0.17 µM and 0 µM (for KCI838PME) and 10 µM, 0.18 µM and 0 µM (for KCI838), respectively **(Figure 6C)** demonstrating the expected rapid clearance of the soluble prodrug. Next, for the antitumor efficacy studies, two groups of 7 mice per group, each bearing bilaterally implanted tumors received 8 daily bolus IV injections of 173 mg/kg KCI838PME in water (treatment group) or water alone (control group). The average weight loss of the treated group from the daily high dose bolus injections of KCI838PME was ≤ 5 percent of the control group weights **(Table 1)** and there were no overt or lasting behavioral changes in the treated group. While the tumor growth rate in the control group was relatively rapid, reaching a tumor burden of nearly 1 g on Day 8, there was a significant reduction in the tumor growth rate in the treated group, which set in after Day 4 and was greater between Day 6 and Day 8. The overall median growth inhibition of 40% observed on Day 8 was highly statistically significant (p=0.0005) but it may be noted that between Day 6 and Day 8, when the maximum tumor growth inhibition occurred, there was a 70% reduction in tumor growth **(Figure 6D)**. Six hours after the last (eighth) bolus injection, the plasma levels of KCI838 and KCI838PME were low as expected, but the active KCI838 alone showed accumulation in the residual tumor samples from the treated mice **(Figure 6E)**.

In the next approach to determining the *in vivo* efficacy of KCI838PME, the prodrug was administered via controlled release using implanted ALZET osmotic pumps. Due to logistical limitations of using implanted ALZET pumps in mice (bulk of the pumps and their capacity), the maximum achievable constant plasma level of KCI838PME was approximately 1 µM over one week, based on the plasma drug levels measured on Day 2 and Day 7 **(Figure 6F)**. Despite this limitation, there was a 62 percent inhibition of tumor growth during the one week of controlled drug release (p=0.0005) **(Figure 6G)**. After Day 7, when the prodrug in the pumps was exhausted, the residual treated tumors resumed the growth rate of the control tumors (measured up to Day 14), demonstrating the expected reversibility of tumor growth inhibition for an AR targeted drug **(Figure 6G)**. Notably, the total daily drug dosing for the controlled release method was less than a fourth of the daily dose used by the bolus injection method **(Figure 6D vs 6G)**.

## Discussion

Following initial surgical or radiological ablation of prostate tumors, disruption of AR signaling continues to be the most effective therapeutic modality for long-term management of PCa for the broader spectrum of patients. This is because variabilities and heterogeneity in tumor proliferation rates, tumor mutation burden, the immune environment of the tumors and defects in homologous recombination repair genes limit the effectiveness of newer cancer treatments such as immune checkpoint inhibitors and PARP inhibitors to small patient groups with metastatic PCa (29, 30, 31). In contrast, the majority of actionable clinical aberrations in advanced PCa were associated with AR (32). It is therefore well recognized that AR-targeted treatments could be strategically expanded to extend the duration of treatment response, particularly by developing drugs that can overcome resistance mechanisms that bypass hormonal activation of AR. Accordingly, there are ongoing efforts to develop small molecules that target functional interactions of AR that reside outside the LBD, including protein-protein interactions of the NTD and the DBD as well as dimerization and protein-DNA interactions of the DBD (33–35). In this space, inhibitors of the interaction of ELK1 with the NTD of AR offer a potential therapy that should not only impact a broad spectrum of AR+ prostate tumors but also, by selectively disrupting a critical growth signaling axis of AR, avoid interference with many non-growth related transcriptional activities of AR required in various normal tissues (27). As discussed below, the present study establishes that the new analog KCI838 meets the conventional criteria for drug-likeness, while recapitulating the molecular target selectivity and PCa-specific biological actions of the parent platform antagonist, KCI807.

The putative binding pocket for KCI807 is rather small, placing a size limit on new pharmaceutical molecules designed to mimic its mechanism of action (28). Accordingly, KCI838 is very similar in overall structure to KCI807, with the exception of having a quinolone scaffold by virtue of a C ring substitution with a nitrogen bearing a trifluoroethyl group in place of the ring oxygen. A quinoline core defines a broad range of currently used and experimental pharmaceuticals for diverse diseases because of its versatility and other desirable features as a pharmacophore (36). The substitution of the N-trifluoroethyl group conferred on KCI838 the ability to completely inhibit growth of the enzalutamide-resistant 22Rv1 PCa cells with earlier onset of the growth inhibition, at a lower dose, compared with KCI807 as well as other KCI838 analogs containing a variety of other substitutions on the C ring nitrogen. KCI838 strongly inhibited growth of CWR22Rv1-AR-EK cells that is exclusively supported by the ligand-independent AR splice variants and was also a better inhibitor of hormone-dependent growth of VCaP and LNCaP PCa cells compared with KCI807 and enzalutamide. Among a variety of cell lines tested, KCI838 only inhibited AR-dependent cell growth. KCI838 inhibited coactivation of ELK1 by AR, both in a promoter-reporter system and in endogenous gene targets. In an orthogonal BRET assay, KCI838 inhibited association of AR and ELK1 *in situ*. The ability of KCI838 to inhibit growth in PCa cells was clearly dependent on its action on AR, as only a two-fold elevation in AR expression using ectopic AR produced the expected shift in dose response of growth inhibition by the compound. Thus, while KCI838 did show more pronounced growth inhibitory effects on a variety of AR+ PCa cell lines compared with KCI807, it did not show any deviation from the mode of action of KCI807 in selectively targeting the AR-ELK1 growth axis.

Scaffold hopping is a common approach in pharmaceutical development to enhance metabolic stability without compromising drug activity (37). An important objective of our study in moving from the flavone scaffold in KCI807 to the quinolone scaffold in KCI838 was to address the problem of self-induced metabolism of KCI807 that limited its ability to inhibit tumor growth *in vivo* in a sustained manner (29). The major enzymes induced by KCI807 in primary human hepatocytes were the monooxygenases CYP1A2 and CYP2B6 and the UDP-glucuronyl transferases UGT1A1 and UGT1A4. While KCI838 was a much weaker inducer of these enzymes compared with KCI807, it should also be noted that the quinolone scaffold is a known inhibitor, rather than a substrate for CYP1A2 (38). Additionally, the relatively poor induction of UGT1A1 and UGT1A4 by KCI838 would suggest that KCI838 is less prone to self-induced metabolism compared with KCI807. The fact that KCI838 did not substantially induce liver enzymes at a ≤ 5 uM dose also suggests that it is unlikely to cause enzyme induction-associated drug-drug interactions (39). KCI838 was also a poorly interacting ligand (substrate or inhibitor) for CYP1A2 and UGT1A1 compared with KCI807, suggesting that it is likely to be more slowly metabolized than KCI807 and also, as a likely inhibitor of CYP1A2, unlikely to cause significant drug-drug interactions associated with liver enzyme inhibition (40).

Considering the hydrophobicity of KCI838, we addressed *in vivo* bioavailability of the compound with a soluble prodrug by converting KCI838 to its 3’-phosphate monoester (KCI838PME). As clinical prostate tumors at all stages are characteristically rich in secreted prostatic acid phosphatase (PAP), the prodrug should be hydrolyzed by the enzyme at the tumor site to release the active drug that should then be efficiently taken up by the tumor cells because of its hydrophobicity. We also confirmed that the PDX tumor used for the *in vivo* testing was positive for (PAP). Despite the rapid systemic clearance of the soluble prodrug KCI838PME administered in the mouse PCa PDX model via daily bolus injection, *in vivo* growth of the tumor was inhibited significantly. The tumor growth inhibition was progressive, with little effect observed initially and maximal effect observed during the later days of treatment. Indeed, the most striking growth inhibition occurred during the last two days of the 8-day treatment. This is presumably related to progressive tumor accumulation of the active hydrophobic drug that was measurable in the tumors at the end of the treatment period.

Although the treated mice did not show obvious signs of toxicity due to KCI838PME, daily administration of the soluble phosphate ester prodrug would not be feasible in patients because of the expected poor oral bioavailability of soluble drugs such as KCI838PME and the susceptibility of KCI838PME to intestinal alkaline phosphatase. Controlled release formulations of soluble drugs are in clinical use; a relevant example is the GnRH agonist leuprolide acetate administered as a subcutaneous hydrogel or as an intramuscular microsphere depot to achieve testosterone suppression to treat advanced PCa with a constant drug release rate lasting several weeks or months. It is therefore logical to consider controlled release of KCI838PME as a treatment modality for advanced PCa that is resistant to current testosterone- or AR-targeted long-term maintenance therapies. Additionally, controlled drug release should avoid any acute or cumulative toxicities caused by relatively high bolus doses. To seek proof-of-principle, we used 1-week release ALZET osmotic pumps implanted in the PDX bearing mice to achieve controlled delivery of KCI838PME into the circulation for evaluation of anti-tumor efficacy of KCI838PME. Although logistical limitations of using ALZET pumps in mice limited dosing of the PDX model with controlled release of KCI838PME, striking tumor growth inhibition was achieved at a mere 1uM plasma level of KCI838PME at less than a fourth of the total daily drug dosage compared with the bolus injection method. The growth inhibition occurred during the 7-day treatment period with the drug and as expected, tumor growth resumed at a rate comparable to control, following depletion of the drug (from Day 8), further confirming the drug effect. As an empirical comparison, leuprolide acetate, has similar solubility compared with KCI838PME (20mg/ml) and its effective dosage when administered through ALZET pumps in mice (41) was similar to the dose used for KCI838PME in this study (32 mg/kg/day for leuprolide vs. 41 mg/kg/day for KCI838PME). These findings offer proof-of-principle for the efficacy and feasibility of treating PCa patients with an appropriate controlled release formulation of KCI838PME.

Steroid hormone targeted cancer therapies, including AR-targeted PCa therapies in current use, are primarily growth-inhibitory, rather than cytotoxic. Cytotoxic therapies, such as PSMA-targeted ^177^Lu radiotherapy for metastatic prostate cancer have several limitations (42, 43). However, tumor regression is indeed known to occur as a secondary effect of prolonged cell cycle arrest in G1 or M phases on cell survival (44, 45). KCI838 should share the non-cytotoxic feature of other AR-targeted drugs, while being effective in a wider spectrum of tumors and additionally, without the need for systemic testosterone suppression. As prostate cancer typically occurs in the older age group, KCI838 should contribute to the goal of extending the life of the patient while potentially offering a relatively better quality of life.

In conclusion, as a rationally designed derivative of the platform antagonist KCI807, the quinolone derivative compound KCI838 shows the expected narrow mechanism of action as an inhibitor of AR-dependent growth of both enzalutamide-sensitive and -resistant PCa cells, with more pronounced effects than KCI807. *In vitro* studies also suggest that KCI838 will meet the criteria for metabolic viability as a pharmaceutical, in contrast to KCI807. Finally, KCI838PME is a promising soluble prodrug of KCI838 as a candidate drug for clinical application in the form of a controlled release formulation to treat advanced PCa. The potential advantages of KCI838PME include 1) Its ability to overcome resistance to current maintenance therapies comprising testosterone suppression and androgen antagonists, 2) A mechanism of action that should obviate the need for castration, 3) Its limited target tissue selectivity for prodrug activation, and 4) Minimization of systemic toxicity via low dose, controlled release.

## Abbreviations

PCa: Prostate cancer
CRPC: Castration resistant prostate cancer
AR: Androgen receptor
AR-V7: Androgen receptor splice variant 7
NTD: Amino-terminal domain
DBD: DNA binding domain
LBD: Ligand binding domain
ETS: Erythroblast transformation specific
ERK: Extracellular signal-regulated kinase
MEK: Mitogen activated protein kinase kinase
BRET: Bioluminescence resonance energy transfer
KCI838PME: Disodium salt of the 3’phosphate monoester of KCI838

## Author Contributions

Claire Soave: Design and performance of most of the *in vitro* experiments and manuscript preparation; Lisa Polin: Design and performance of animal model studies; Charles Ducker: Development, design and performance of BRET assays; Vy Ong: Statistical analyses; Seongho Kim: Statistical analyses; Luke Pardy: Modeling and synthetic chemistry; Jing Li: Direction of pharmacokinetic analyses; Xun Bao: Development and conduct of LC/MS assays; Yanfang Huang: Technical support; Peter Shaw: Sharing expertise on ELK1, development of BRET assays; Rahul Khupse: Sharing expertise in synthetic chemistry; Manohar Ratnam: Overall conception and direction of all aspects of the project, small molecule design, manuscript preparation.

## Acknowledgements

This work was supported in part by the U.S. Department of Defense [Grant W81XWH-17-1-0242] (to M.R.) and [Grant W81XWH-17-1-0243] (to P.E.S.) and National Institutes of Health National Cancer Institute [Grant 5T32-CA009531-29] (to C.S.).

## Supplemental Figures

**Supplemental Figure 1.**
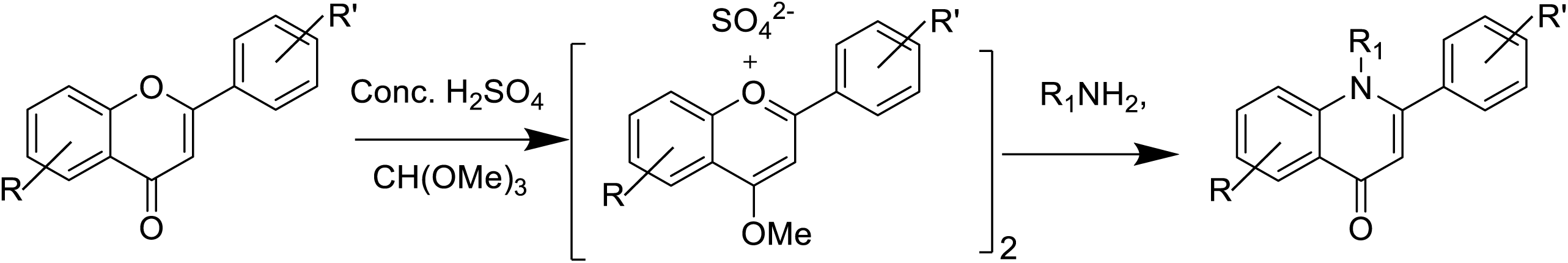

**Supplemental Figure 2.**
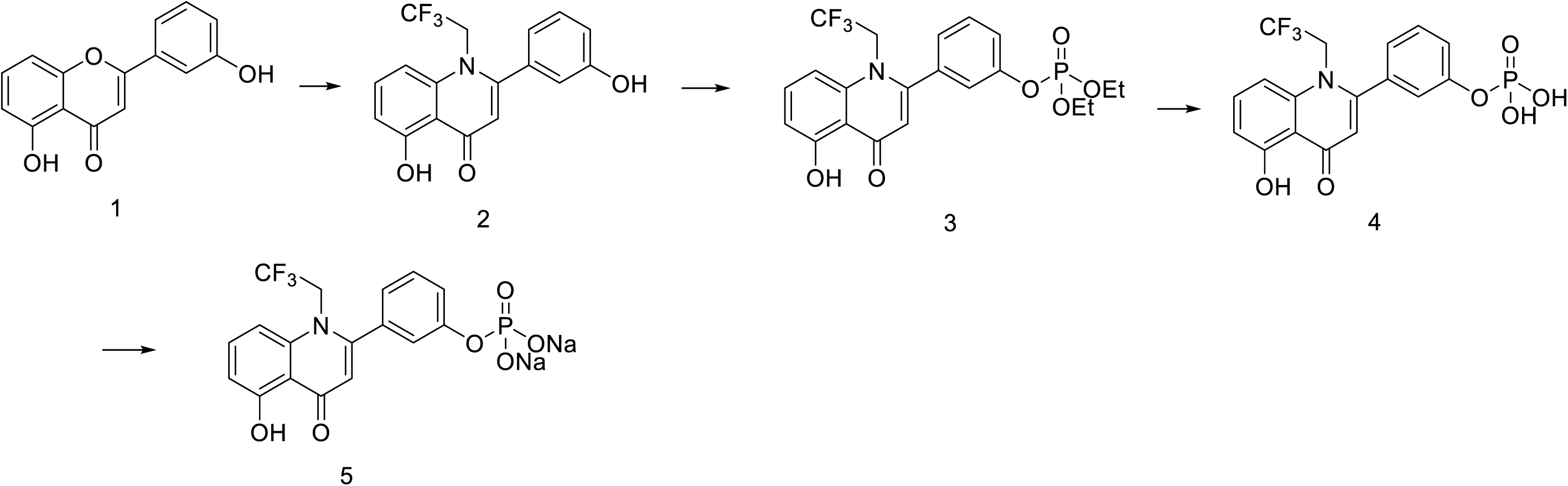

## References

1. Massard C, Fizazi K. Targeting continued androgen receptor signaling in prostate cancer. Clin Cancer Res. 2011;17(12):3876–83.

2. Ryan CJ, Tindall DJ. Androgen receptor rediscovered: the new biology and targeting the androgen receptor therapeutically. J Clin Oncol. 2011;29(27):3651–8.

3. Dunn MW, Kazer MW. Prostate cancer overview. Semin Oncol Nurs. 2011;27(4):241–50.

4. Loblaw DA, Virgo KS, Nam R, Somerfield MR, Ben-Josef E, Mendelson DS, et al. Initial hormonal management of androgen-sensitive metastatic, recurrent, or progressive prostate cancer: 2006 update of an American Society of Clinical Oncology practice guideline. J Clin Oncol. 2007;25(12):1596–605.

5. Rice MA, Malhotra SV, Stoyanova T. Second-Generation Antiandrogens: From Discovery to Standard of Care in Castration Resistant Prostate Cancer. Frontiers in Oncology. 2019;Volume 9 - 2019.

6. Yin L, Hu Q. CYP17 inhibitors--abiraterone, C17,20-lyase inhibitors and multi-targeting agents. Nat Rev Urol. 2014;11(1):32–42.

7. Lamont KR, Tindall DJ. Minireview: Alternative activation pathways for the androgen receptor in prostate cancer. Mol Endocrinol. 2011;25(6):897–907.

8. Chan SC, Li Y, Dehm SM. Androgen receptor splice variants activate androgen receptor target genes and support aberrant prostate cancer cell growth independent of canonical androgen receptor nuclear localization signal. J Biol Chem. 2012;287(23):19736–49.

9. Cao B, Qi Y, Zhang G, Xu D, Zhan Y, Alvarez X, et al. Androgen receptor splice variants activating the full-length receptor in mediating resistance to androgen-directed therapy. Oncotarget. 2014;5(6):1646–56.

10. Sprenger CC, Plymate SR. The link between androgen receptor splice variants and castration-resistant prostate cancer. Horm Cancer. 2014;5(4):207–17.

11. Li Y, Chan SC, Brand LJ, Hwang TH, Silverstein KA, Dehm SM. Androgen receptor splice variants mediate enzalutamide resistance in castration-resistant prostate cancer cell lines. Cancer Res. 2013;73(2):483–9.

12. Sun S, Sprenger CC, Vessella RL, Haugk K, Soriano K, Mostaghel EA, et al. Castration resistance in human prostate cancer is conferred by a frequently occurring androgen receptor splice variant. J Clin Invest. 2010;120(8):2715–30.

13. Hu R, Lu C, Mostaghel EA, Yegnasubramanian S, Gurel M, Tannahill C, et al. Distinct transcriptional programs mediated by the ligand-dependent full-length androgen receptor and its splice variants in castration-resistant prostate cancer. Cancer Res. 2012;72(14):3457–62.

14. Nyquist MD, Li Y, Hwang TH, Manlove LS, Vessella RL, Silverstein KAT, et al. TALEN-engineered AR gene rearrangements reveal endocrine uncoupling of androgen receptor in prostate cancer. Proceedings of the National Academy of Sciences. 2013;110(43):17492–7.

15. Henzler C, Li Y, Yang R, McBride T, Ho Y, Sprenger C, et al. Truncation and constitutive activation of the androgen receptor by diverse genomic rearrangements in prostate cancer. Nature Communications. 2016;7(1):13668.

16. Kounatidou E, Nakjang S, McCracken SRC, Dehm SM, Robson CN, Jones D, et al. A novel CRISPR-engineered prostate cancer cell line defines the AR-V transcriptome and identifies PARP inhibitor sensitivities. Nucleic Acids Res. 2019;47(11):5634–47.

17. Myklak K, Wilson S. An update on the changing indications for androgen deprivation therapy for prostate cancer. Prostate Cancer. 2011;2011:419174.

18. Holzbeierlein JM, McLaughlin MD, Thrasher JB. Complications of androgen deprivation therapy for prostate cancer. Curr Opin Urol. 2004;14(3):177–83.

19. Shaw PE, Saxton J. Ternary complex factors: prime nuclear targets for mitogen-activated protein kinases. Int J Biochem Cell Biol. 2003;35(8):1210–26.

20. Gille H, Sharrocks AD, Shaw PE. Phosphorylation of transcription factor p62TCF by MAP kinase stimulates ternary complex formation at c-fos promoter. Nature. 1992;358(6385):414–7.

21. Zhang HM, Li L, Papadopoulou N, Hodgson G, Evans E, Galbraith M, et al. Mitogen-induced recruitment of ERK and MSK to SRE promoter complexes by ternary complex factor Elk-1. Nucleic Acids Res. 2008;36(8):2594–607.

22. Yu J, Yu J, Mani RS, Cao Q, Brenner CJ, Cao X, et al. An integrated network of androgen receptor, polycomb, and TMPRSS2-ERG gene fusions in prostate cancer progression. Cancer Cell. 2010;17(5):443 –54.

23. Patki M, Chari V, Sivakumaran S, Gonit M, Trumbly R, Ratnam M. The ETS domain transcription factor ELK1 directs a critical component of growth signaling by the androgen receptor in prostate cancer cells. J Biol Chem. 2013;288(16):11047–65.

24. Rosati R, Patki M, Chari V, Dakshnamurthy S, McFall T, Saxton J, et al. The Amino-terminal Domain of the Androgen Receptor Co-opts Extracellular Signal-regulated Kinase (ERK) Docking Sites in ELK1 Protein to Induce Sustained Gene Activation That Supports Prostate Cancer Cell Growth. Journal of Biological Chemistry. 2016;291(50):25983–+.

25. Soave C, Ducker C, Kim S, Strahl T, Rosati R, Huang Y, et al. Identification of ELK1 interacting peptide segments in the androgen receptor. Biochem J. 2022;479(14):1519–31.

26. Pardy L, Rosati R, Soave C, Huang Y, Kim S, Ratnam M. The ternary complex factor protein ELK1 is an independent prognosticator of disease recurrence in prostate cancer. Prostate. 2020;80(2):198–208.

27. Rosati R, Polin L, Ducker C, Li J, Bao X, Selvakumar D, et al. Strategy for Tumor-Selective Disruption of Androgen Receptor Function in the Spectrum of Prostate Cancer. Clin Cancer Res. 2018;24(24):6509 –22.

28. Soave C, Ducker C, Islam N, Kim S, Yurgelevic S, Nicely NI, et al. The Small Molecule Antagonist KCI807 Disrupts Association of the Amino-Terminal Domain of the Androgen Receptor with ELK1 by Modulating the Adjacent DNA Binding Domain. Mol Pharmacol. 2023;103(4):211–20.

29. Liu D, Wang L, Guo Y. Advances in and prospects of immunotherapy for prostate cancer. Cancer Lett. 2024 Oct 1;601:217155. doi: 10.1016/j.canlet.2024.217155. Epub 2024 Aug 8. PMID: 39127338.

30. Claps M, Mennitto A, Guadalupi V, Sepe P, Stellato M, Zattarin E, et al. Immune-checkpoint inhibitors and metastatic prostate cancer therapy: Learning by making mistakes. Cancer Treat Rev. 2020;88:102057.

31. Teyssonneau D, Margot H, Cabart M, Anonnay M, Sargos P, Vuong NS, et al. Prostate cancer and PARP inhibitors: progress and challenges. J Hematol Oncol. 2021;14(1):51.

32. Robinson D, Van Allen EM, Wu YM, Schultz N, Lonigro RJ, Mosquera JM, et al. Integrative clinical genomics of advanced prostate cancer. Cell. 2015;161(5):1215–28.

33. Tan MH, Li J, Xu HE, Melcher K, Yong EL. Androgen receptor: structure, role in prostate cancer and drug discovery. Acta Pharmacol Sin. 2015;36(1):3–23.

34. Monaghan AE, McEwan IJ. A sting in the tail: the N-terminal domain of the androgen receptor as a drug target. Asian Journal of Andrology. 2016;18(5):687–94.

35. Monaghan AE, Porter A, Hunter I, Morrison A, McElroy SP, McEwan IJ. Development of a High-Throughput Screening Assay for Small-Molecule Inhibitors of Androgen Receptor Splice Variants. Assay Drug Dev Technol. 2022;20(3):111–24.

36. Yadav P, Shah K. Quinolines, a perpetual, multipurpose scaffold in medicinal chemistry. Bioorg Chem. 2021;109:104639.

37. Lazzara PR, Moore TW. Scaffold-hopping as a strategy to address metabolic liabilities of aromatic compounds. RSC Med Chem. 2020;11(1):18–29.

38. Fuhr U, Strobl G, Manaut F, Anders EM, Sörgel F, Lopez-de-Brinas E, et al. Quinolone antibacterial agents: relationship between structure and in vitro inhibition of the human cytochrome P450 isoform CYP1A2. Mol Pharmacol. 1993;43(2):191–9.

39. Bünning P. Drug–Drug Interaction: Enzyme Induction. In: Vogel HG, Maas J, Hock FJ, Mayer D, editors. Drug Discovery and Evaluation: Safety and Pharmacokinetic Assays. Berlin, Heidelberg: Springer Berlin Heidelberg; 2013. p. 975–87.

40. Dudda A, Kuerzel GU. Drug–Drug Interaction: Enzyme Inhibition. In: Vogel HG, Maas J, Hock FJ, Mayer D, editors. Drug Discovery and Evaluation: Safety and Pharmacokinetic Assays. Berlin, Heidelberg: Springer Berlin Heidelberg; 2013. p. 989–1004.

41. Xi L, Kraskauskas D, Muniyan S, Batra SK, Kukreja RC. Androgen-deprivation therapy with leuprolide increases abdominal adiposity without causing cardiac dysfunction in middle-aged male mice: effect of sildenafil. American Journal of Physiology-Regulatory, Integrative and Comparative Physiology. 2023;324(4):R589–R600.

42. Sartor O, Herrmann K. Prostate Cancer Treatment: (177)Lu-PSMA-617 Considerations, Concepts, and Limitations. J Nucl Med. 2022;63(6):823–9.

43. Sadaghiani MS, Sheikhbahaei S, Werner RA, Pienta KJ, Pomper MG, Solnes LB, et al. A Systematic Review and Meta-analysis of the Effectiveness and Toxicities of Lutetium-177-labeled Prostate-specific Membrane Antigen-targeted Radioligand Therapy in Metastatic Castration-Resistant Prostate Cancer. Eur Urol. 2021;80(1):82–94.

44. Crozier L, Foy R, Mouery BL, Whitaker RH, Corno A, Spanos C, et al. CDK4/6 inhibitors induce replication stress to cause long-term cell cycle withdrawal. Embo j. 2022;41(6):e108599.

45. Orth JD, Loewer A, Lahav G, Mitchison TJ. Prolonged mitotic arrest triggers partial activation of apoptosis, resulting in DNA damage and p53 induction. Mol Biol Cell. 2012;23(4):567–76.

